# Noradrenergic *locus coeruleus* activity functionally partitions NREM sleep to gatekeep the NREM-REM sleep cycle

**DOI:** 10.1101/2023.05.20.541586

**Authors:** Alejandro Osorio-Forero, Georgios Foustoukos, Romain Cardis, Najma Cherrad, Christiane Devenoges, Laura M.J. Fernandez, Anita Lüthi

**Affiliations:** Department of Fundamental Neurosciences, University of Lausanne, Rue du Bugnon 9, CH-1005 Lausanne, Switzerland; 2 Department of Sleep and Cognition. Netherlands Institute for Neuroscience, Meibergdreef 47, 1105 BA Amsterdam, The Netherlands

**Keywords:** ultradian sleep cycle, arousability, sleep homeostasis, REMS restriction, automated REMS deprivation, microarousal, noradrenaline, norepinephrine, infra-slow time scale, optogenetics, K-complex, sleep disorders

## Abstract

The noradrenergic *locus coeruleus* (LC) is vital for brain states underlying wakefulness, whereas its roles for sleep remain uncertain. Combining mouse sleep-wake monitoring, behavioral manipulations, LC fiber photometry and closed-loop optogenetics, we found that LC neuronal activity partitioned non-rapid-eye-movement sleep (NREMS) into alternating brain and autonomic states that rule the NREMS-REMS cycle. High LC activity levels generated an autonomic-subcortical arousal state that facilitated cortical microarousals, while low levels were obligatory for REMS entries. Timed optogenetic LC inhibition revealed that this functional alternation set the duration of the NREMS-REMS cycle by ruling REMS entries during undisturbed sleep and when pressure for REMS was high. A stimulus-enriched, stress-promoting wakefulness increased high LC activity levels at the expense of low ones in subsequent NREMS, fragmenting NREMS through microarousals and delaying REMS onset. We conclude that LC activity fluctuations gatekeep the NREM-REMS cycle over recurrent infraslow intervals, but they also convey vulnerability to adverse wake experiences.

## Introduction

The pontine brainstem *locus coeruleus* (LC) is a wake-promoting brain area that constitutes the principal source of noradrenaline (NA) for forebrain circuits and alertness-modulating nuclei^1^. LC neuronal activity levels grade with the level of wakefulness^2^ and with physiological signatures of cortical^3^ and autonomic arousal^4,5^, while sleep-promoting brain areas are inhibited^6,7^. Elevated LC activity is implicated in the cognitive and autonomic manifestations of states of hyperarousal during stress-and anxiety-provoking experiences^1,8^. Conversely, clinically important sedatives inhibit LC^9^. Together, this illustrates LC’s broad implication in regulating arousal levels in states of wakefulness.

Many noradrenergic LC neurons remain active in sleep^7,10–14^, during which they increase arousability^15,16^, in line with LC’s wake-promoting actions^1,3^. During non-rapid-eye-movement sleep (NREMS) in mouse, activity levels of LC neuron populations fluctuate on an infraslow time scale (30 – 50 s, 0.02 Hz), generating varying levels of free NA in the forebrain^17–19^. For example, in thalamus, NA fluctuations recurrently depolarize thalamocortical and thalamic reticular neurons. This depolarization suppresses the capacity of thalamic circuits to generate sleep spindles, known as NREMS EEG hallmarks in the 10 – 15 Hz frequency range^17^ that shield NREMS from sensory disruption^20^. The LC-induced thalamic activation hence promotes auditory and somatosensory arousability^16,21,22^. Spontaneous microarousals (MAs) that are typical for mammalian NREMS also coincide preferentially with moments of low spindle density^18,21^. When reaching high levels, LC activity fluctuations generate brain states with high arousability, suggesting that they could be primarily sleep-disruptive. Alternatively, its activity fluctuations could have a broader relevance for the dynamics of sleep. For example, for sleep to take its natural progression across NREMS and REMS, it is necessary that global brain states be driven across transitory, less stable periods. In human sleep, signatures of sleep ‘instability’ and elevated arousability have been described^23^, yet neither the underlying neural mechanisms driving these nor the associated brain states are known.

A low level of LC unit activity has further been associated with REMS^10–12^. This prompted the first circuit-based theoretical models for the mammalian NREM-REM sleep cycle, in which the LC was integrated as a REM-off area^24,25^. Indeed, optogenetic activation of NA-synthetizing LC neurons can increase the time spent in NREMS at the expense of REMS, whereas inhibition can have opposite effects^15,17–19^. Also, widely used sedatives that interfere with NA signaling modify REMS in humans^26,27^. Furthermore, the level of LC activity in wakefulness negatively associated with human REMS quality^28^. Therefore, a balanced assessment of LC actions REMS-regulatory and REMS-disruptive actions is desirable and is indeed pressing as its implication in human sleep becomes increasingly recognized^29,30^.

Here, we show that LC rules the time scales over which the NREMS-REMS cycle evolves, setting the moments when transitions between NREMS and REMS are allowed, and when not. This ruling was enabled through the alternation of LC activity fluctuations between high and low levels that induced distinct brain states characterized by a unique profile of autonomic, subcortical and cortical arousal levels. High levels of LC activity fluctuations preferentially lead to MAs but suppressed REMS entries. REMS entries took place once LC activity decayed to low levels, either spontaneously or through optogenetic inhibition. We further found that the infraslow temporal alternation between high and low LC activity levels regulated the duration of the NREMS-REMS cycle and restrained it to a minimum of ∼50 s during REMS restriction (REMS-R). After a stimulus-enriched wake experience, LC activity fluctuations became unbalanced and persisted at high levels, which caused more arousals and delayed REMS. The LC is hence a neural mechanism that gatekeeps the NREMS-REMS-cycle duration, but that can generate cycle disruptions as a result of preceding wake experience.

## Results

### High LC activity levels during NREMS promoted brain and autonomic activation that facilitates arousability

To monitor the activity of LC neurons during unperturbed sleep, we conditionally expressed the genetically encoded Ca^2+^ sensor jGCaMP8s in dopamine-β-hydroxylase-(DBH)-Cre mice, previously shown to drive the expression of viral constructs in NA-synthetizing neurons with a high degree of specificity (> 80 %, see also **Extended Data Fig. 1a**)^17,31^. We then implanted animals with EEG/EMG electrodes together with an optic fiber stub over the LC for combined sleep and fiber photometric recordings of LC neuronal activity (**Fig. 1a, b**). Local field potential (LFP) electrodes were additionally implanted in the primary somatosensory S1 cortex and the hippocampal CA1 area. The EEG signals served to score the major vigilance states Wake, NREMS and REMS and the MAs based on standard polysomnography^17,21,22,32^. The S1 LFP recordings were used to monitor the local thalamocortical dynamics of power bands characteristic for NREMS and the CA1 LFP recordings to define the onset of REMS (see Methods).

**Figure 1.**
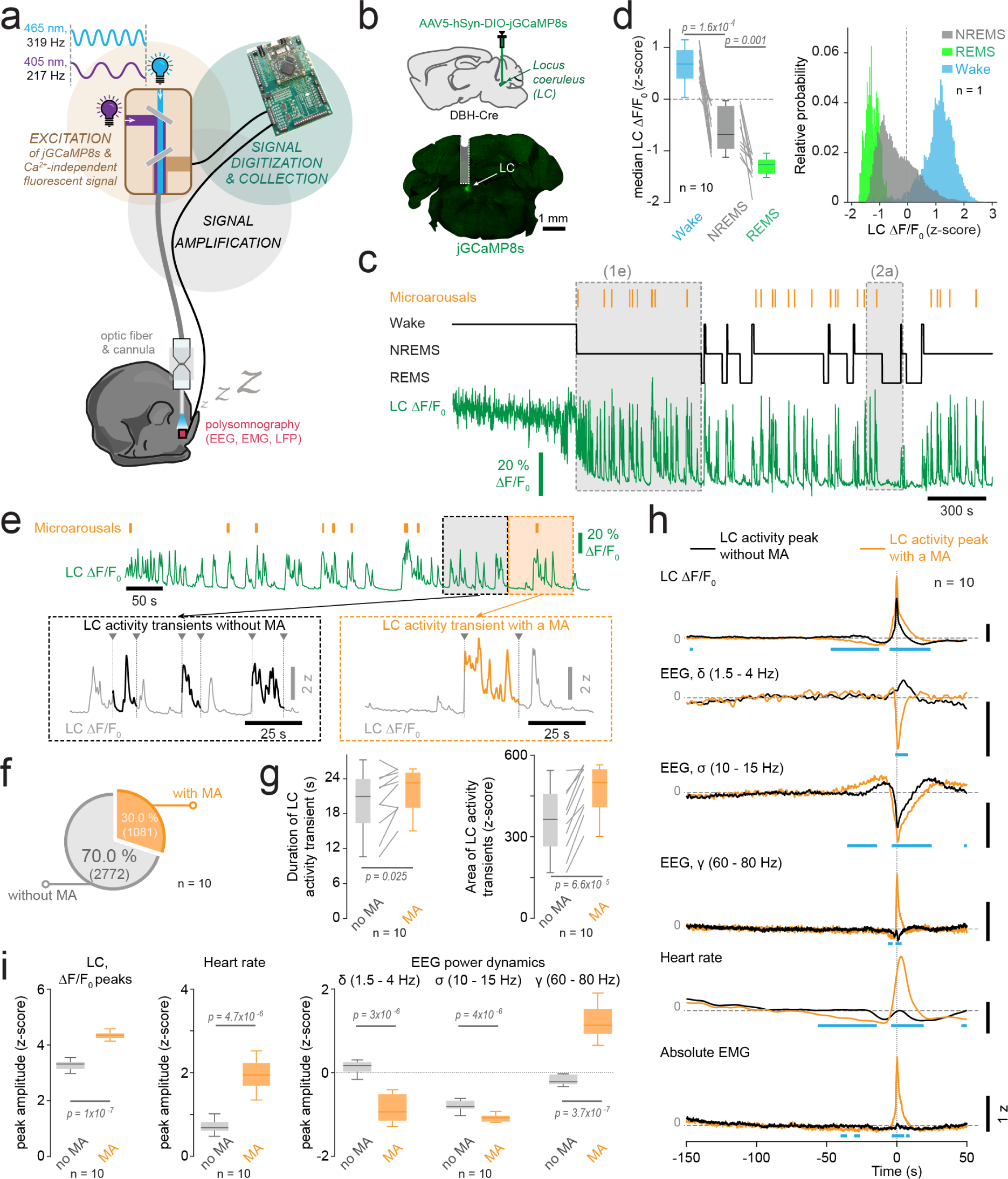
*Locus coeruleus* (LC) activity patterns across wakefulness, NREMS and REMS. a) Experimental scheme for sleep recordings in combination with fiber photometry. b) Schematic of viral injection into a dopamine-β-hydroxylase (DBH)-Cre mouse prior to sleep/fiber photometric implantations and example *post hoc* histological verification of jGCaMP8s expression and optic fiber location. c) Example sleep hypnogram (black) and jGCaMP8s fluorescent signal (green) recorded from LC (in ΔF/F_0_). Orange vertical lines: microarousals (MAs). Grey rectangles: trace portions that are expanded in other figure panels (as indicated). d) Left, Box-and-whisker plot of mean LC fluorescence levels for wake, NREMS and REMS for Zeitgeber times (ZT) ZT0-ZT2 for n = 10 animals, with grey lines connecting values for individual animals; one-way repeated-measures ANOVA with *post hoc* t-tests. Bonferroni-corrected p = 0.025. Right, all-points-histogram for z-scored fluorescence across wake, NREMS and REMS for a representative animal. e) Example trace expanded from panel c for NREMS. Orange vertical lines show MAs. Colored dotted rectangles indicate two trace portions expanded below. For these expanded portions, thick traces show LC activity transients without MAs (left) and with a MA (right). The on-and offset of these transients is marked with arrowheads connected to vertical lines. For further details, please see Extended Data Fig. 2. f) Pie chart for proportion of LC activity transients without and with MAs in % for n = 10 animals for ZT0-ZT2. Numbers in parentheses include total counts of LC activity transients. g) Box-and-whisker plots for duration (left) and area (right) of LC activity transients identified as illustrated in panel e for n = 10 animals. Grey lines connect mean values from individual animals. h) Spectral analysis of EEG signals corresponding to LC activity peaks that were classified based on whether they occurred without a MA (black) or with a MA (orange). Corresponding power dynamics for the delta (δ), sigma (σ) and gamma (γ) frequency bands are aligned vertically, together with heart rate and absolute EMG. Traces denote means across n = 10 animals. Blue bars denote significance p values as calculated by false discovery rates. i) Quantification of mean values from traces shown in h for peak values in the time interval 0-1s. All p values were calculated from paired *t*-tests. For power analyses, the Bonferroni-corrected p was 0.017. See **Supplementary Table 1** for detailed statistical information.

Across the 12-h light phase (ZT0-ZT12), the major resting phase of mice, the photometrically recorded Ca^2+^ activity of noradrenergic LC populations (‘LC activity’) varied characteristically with states of wake, NREMS and REMS, consistent with recent studies (**Fig. 1c**)^18,19^. LC activity was elevated during wakefulness, but it declined in NREMS to a lower baseline on top of which a fluctuating pattern appeared. In 10 animals with a dynamic range of relative fluorescence changes between 16.3 – 95.3 % (mean 41.6 ± 9.3 %) in the first 2 h of the light phase (ZT0-ZT2) and a histologically confirmed fiber positioning (**Fig. 1b**), LC fluctuations were present throughout NREMS bouts but were absent during REMS (**Fig. 1c**). Mean LC activity levels in ZT0-ZT2 declined progressively from wakefulness to NREMS and REMS (**Fig. 1d**) (for statistical details in all figures, see **Supplementary Table 1**). However, activity levels in NREMS varied widely, with 56.5 ± 11.4 % overlapping with wakefulness (calculated up to the mean - 2 standard deviations of wake levels; **Fig. 1d, Extended Data Fig. 1b**). In mouse NREMS, the collective neuronal activity levels of the noradrenergic LC could thus go as high as in some periods of wakefulness.

Elevated LC activity promotes arousability from NREMS^16^, but the brain states underlying this arousal-promoting action have not been systematically examined. The LC projects cortically and subcortically, including into autonomic output centers^1,7^, suggesting that LC regulates arousability through coordinating brain-bodily arousal levels. We first confirmed that LC activity fluctuations generated robust variations of free NA levels in forebrain. Dual fiber photometric measures based on jGCaMP8s fluorescence in LC and on GRAB_NE1h_ fluorescence^33^ in thalamus confirmed phase-locked variations in both signals (cross-correlation coeff. > 0.5) (**Extended Data Fig. 1c, d**). Furthermore, high LC activity levels coincided with power declines in a NREMS-specific frequency band, the spindle-containing σ (10 – 15 Hz) power band, whereas σ power rose during low LC activity levels (**Extended Data Fig. 1e1**)^17–20^. LC activity and σ power were anticorrelated with a time lag of 2.1 ± 0.5 s (n = 10) and showed side peaks at 48.3 ± 2.2 s (**Extended Data Fig. 1e2**, range 38.5 – 58.1 s, measured in n = 6/10 animals in which these side peaks were clearly detectable). Therefore, in line with recent findings^17^, LC activity transients generated robust NA release that activated thalamus through depolarization and suppressed sleep spindles.

We next isolated LC activity transients and studied the associated brain states together with heart rate as a measure of autonomic activation (**Fig. 1e, Extended Data Fig. 2**). We noted that 30.0 % of LC activity transients (1,081 of 3,853 detected cases in n = 10 animals) were accompanied by a MA (**Fig. 1f**), recognizable by an EEG desynchronization and a concomitant EMG activity lasting ≤12 s^21,32,34^. LC activity transients during MA lasted longer and were larger than when there was no MA, suggesting greater overall noradrenergic output (**Fig. 1g**, n = 10). We hence analyzed non-MA-associated LC activity transients separately from MA-associated ones (**Fig. 1h**, n = 10). These non-MA-associated transients were accompanied by a decrease in σ (10 – 15 Hz) power and an increased heart rate, reflecting a coordinated thalamic and cardiac activation during continuous NREMS. MA-associated LC transients similarly altered sigma power and heart rate, yet with larger effect size (**Fig. 1i**). However, the two types of LC activity transients coupled oppositely to low-frequency power changes. Specifically, power in the δ range (1.5 – 4 Hz) showed a small positive increase for non-MA-associated LC activity transients (p = 0.035, paired t-test, effect size D = −1.02, n = 10; **Fig. 1h, i**), which indicates a period of cortical deactivation or recurrent ‘downstates’^35^. The MA-associated transient showed instead a pronounced decrease in δ power, which reflects a cortical activation.

Opposite dynamics were also obtained for the γ frequency band (60 – 80 Hz), another correlate of cortical activation^36^. In the case of MAs, there were γ power increases that coincided with EMG activity, a MA-defining feature. The EEG spectral dynamics across δ, σ and γ bands could be similarly reproduced for the LFP recordings in the S1 area, which indicates that LC-associated brain state dynamics involved local thalamocortical circuits (**Extended Data Fig. 3a, b**). Point-by-point analyses strongly suggested that all EEG and LFP spectral power changes and the heart rate accelerations were likely induced by LC activity starting less than 50 s from the peak rather than by divergent EEG or LFP dynamics starting before (> 50 s before the peak) (**Fig. 1h, Extended Data Fig. 3a**). These analyses demonstrate that LC activity transients during NREMS coordinate a previously undescribed brain-bodily state, characterized by a combined cardiac and thalamic activation that is dissociated from cortical activation. During this state, arousability was enhanced because MAs appeared preferentially^18,21^, which caused cortical activation.

### Low LC activity levels are required for transitions to REMS

Low LC activity levels have been previously associated with NREMS-to-REMS transitions (‘REMS entries’), but the causal role of the natural activity troughs for REMS entries has not been tested directly. As reported^18,19^, we noted that these troughs were associated with REMS entries (n = 9) (**Fig. 2a**). We next optogenetically modified activity troughs specifically and bidirectionally in DBH-Cre animals expressing the excitatory opsin channelrhodopsin-2 (ChR2) or the inhibitory opsin Jaws in LC neurons^17^. We tested whether REMS entries no longer occurred when LC was activated at moments of the troughs. Conversely, we inhibited LC when it normally reached high levels. We tracked LC fluctuations on-line through monitoring the σ power dynamics in the S1 LFP. We had previously demonstrated that such monitoring allows to reliably distinguish between high and low LC activity levels^17^.

**Figure 2.**
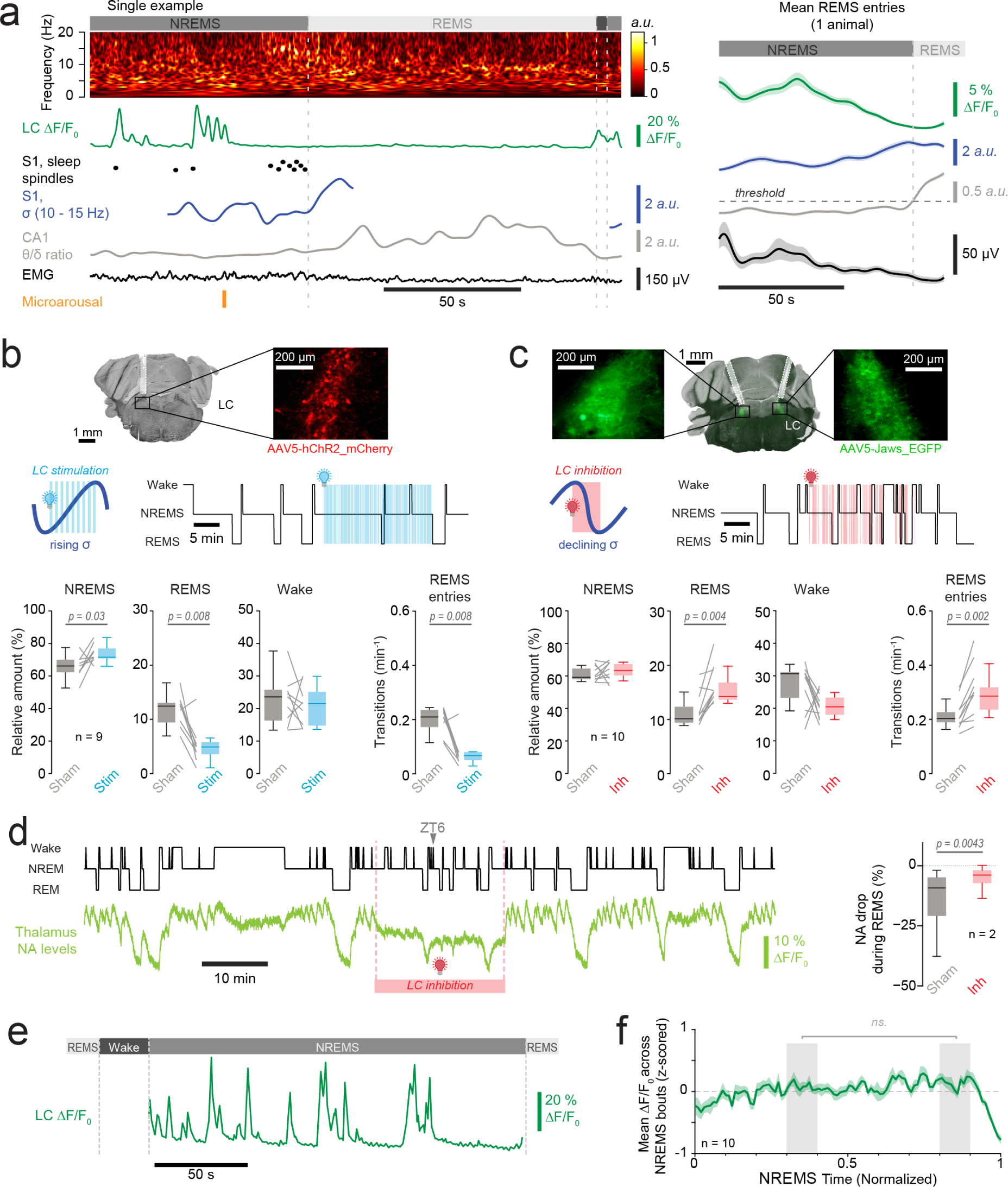
Association of LC activity patterns during NREMS with arousal promotion and REMS entries. a) Left, Example recording of LC fluorescence during a NREMS-to-REMS transition (‘REMS entry’, expanded from Fig. 1c). Same signals shown as in a, with in addition CA1 θ/δ ratio and EMG. Right, Mean dynamics of these signals for the same animal. b) Optogenetic stimulation of LC. Top, Histological verification of fiber positioning and viral transfection. Middle, Schematic of the light stimulation protocol based on closed-loop feedback analysis of σ power. Blue vertical lines show timing of optogenetic stimulation. Example portion of hypnogram is shown on the right. Bottom, Box-and-whisker-plot quantifications were calculated for the 20-min light *vs* sham (LED-off) stimulation periods for n = 9 animals. Grey lines, paired datasets per animal. Wilcoxon signed-rank tests. c) As a), for optogenetic inhibition of LC. d) Example dual fiber photometric recording from an animal that expressed Jaws in LC and GRAB_NE1h_ in somatosensory thalamus. Portion underlain with orange square indicates period of LC inhibition. Quantification was done in n = 2 animals for 45 REMS bouts each in control and in light illumination. REMS bouts in both conditions were selected for equal durations before evaluating GRAB_NE1h_ fluorescence decreases. e) Example trace for LC fluorescence during a NREMS bout. f) Mean LC fluorescence activity across time-normalized NREMS bouts. Shaded areas, data used for statistical analysis. Paired t-test. See **Supplementary Table 1** for detailed statistical information.

We optogenetically activated LC in ChR2-expressing animals at moments of rising σ power (corresponding to low LC activity levels, see also **Extended Data Fig. 1e1**) in the first 20 min of every h during ZT1-ZT9, with corresponding sham stimulations (LEDs turned-off) in separate sessions (**Fig. 2b**)^17^. This reduced the time spent in REMS and the frequency of REMS entries compared to sham periods, but did not alter the density of MAs (0.58 ± 0.08 / min vs 0.62 ± 0.14 /min, Wilcoxon’s signed rank test; p = 0.3; n = 9). Conversely, we expressed the orange light-activated chloride pump ‘Jaws’ in DBH-Cre animals and inhibited LC at moments when it was naturally high. We verified the efficacy of this inhibition in 2 DBH-Cre animals expressing Jaws in the LC and GRAB_NE1h_ in the thalamus. Jaws-inhibition of LC (with continuous light exposure^17^) decreased NA levels in baseline and reduced the extent of NA decline during REMS (**Fig. 2c**). When we next inhibited LC as soon as we detected a decline in σ power, we specifically increased REMS time and the frequency of REMS entries (**Fig. 2d**). Therefore, an optogenetically imposed decrease in natural LC activity, such as it occurs in troughs, was permissive for REMS entries, while an increase in LC activity suppressed such entries.

The propensity to enter REMS increases linearly with the time spent in NREMS, which reflects its homeostatic regulation^37^. Furthermore, several REMS-regulating brain areas innervate the LC^38–41^. Therefore, it was important to test whether LC activity fluctuations show a homeostatic downregulation in the course of NREMS that would eventually permit REMS entries. Contrary to this possibility, mean LC activity levels across NREMS bouts remained stable until just before REMS onset, arguing against homeostatic regulation (**Fig. 2e**). These experiments establish that LC activity troughs, whether naturally occurring or optogenetically promoted, generated windows of opportunity for REMS entries throughout the major time spent in NREMS.

### REMS restriction shows that LC activity fluctuations ruled the timing of REMS entries

The alternation between high and low LC activity levels takes place on infraslow time intervals. Therefore, the time it takes for LC activity to decay from high to low levels could determine the duration of NREMS before a REMS entry becomes possible. Clarifying this possibility would indicate that there is an ultradian time interval that models of the NREMS-REMS cycle would need to integrate^42^. To test this, we sought to increase the frequency of REMS entries while monitoring their timing relative to on-going LC activity. We developed a REMS restriction (REMS-R) method in mice designed to maximally reduce time spent in REMS while preserving NREMS time. We used vibrating motors taped to the animals’ headstage to interrupt REMS in closed-loop feedback (**Fig. 3a**)^21^. Starting at ZT0, these motors vibrated after 2 s of automatically detected REMS. When interrupting every detected entry into REMS over a 6-h period from ZT0-ZT6, we observed a strong increase in attempts to enter REMS (**Fig. 3b**). No such increase was observed in baseline recordings or in yoked control animals that were exposed to vibration signals derived from the animal in the neighboring cage. The REMS-R further preserved homeostatic regulation of NREMS and induced a rebound REMS (**Extended Data Fig. 4**).

**Figure 3.**
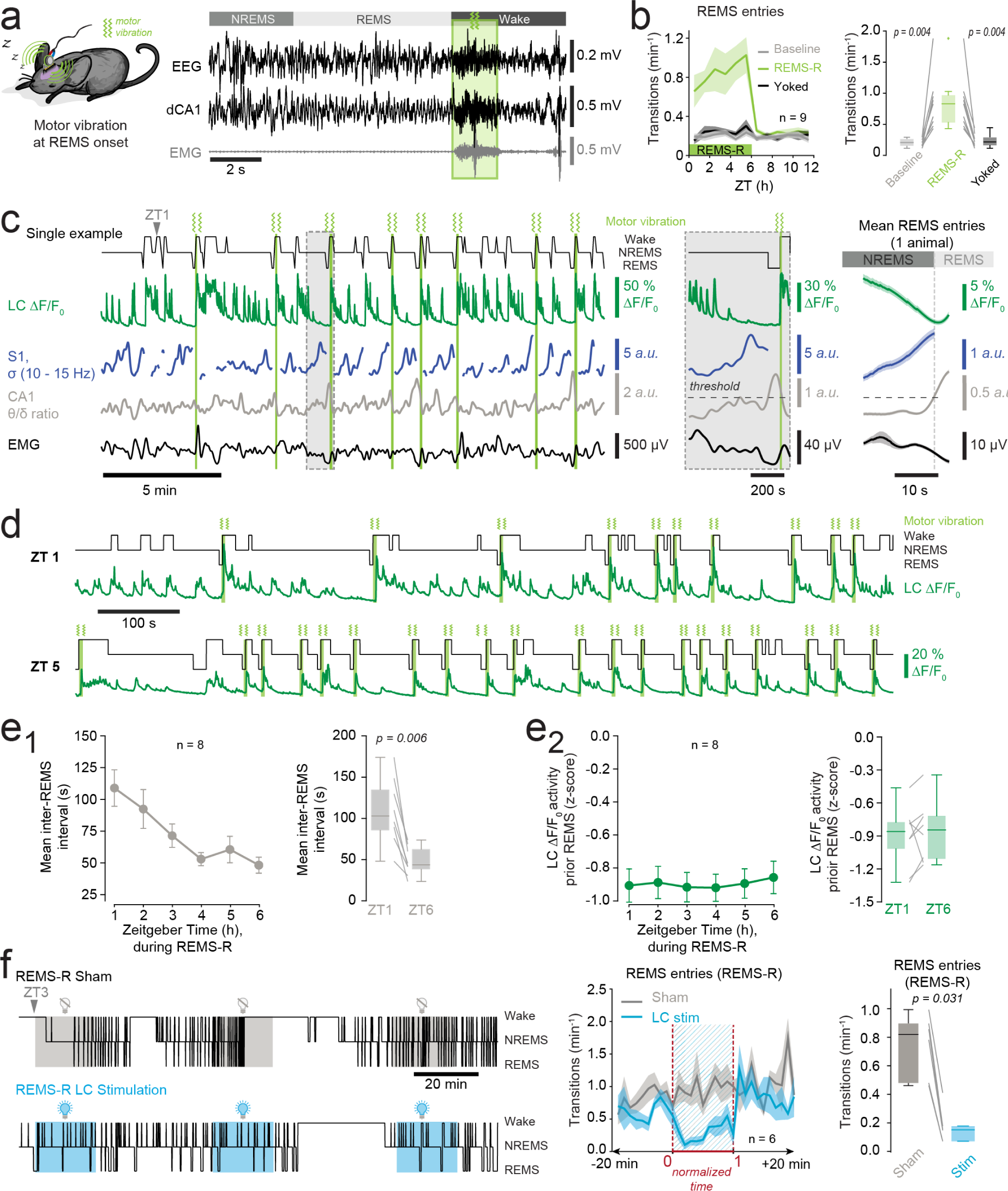
REMS restriction (REMS-R) unravels a minimal duration of the NREMS-REMS cycle related to LC activity decline. a) Schematic of REMS-R implementation and representative recording showing (green rectangle with motor vibration symbol). b) Frequency of REMS entries for n = 9 animals subjected to REMS-R from ZT0-ZT6, to yoked conditions (animals given vibrations timed to REMS from their neighbors) and to undisturbed conditions (baseline), followed by 6 h of recovery. To the right, box-and-whisker plot quantifications, with grey lines showing paired data for individual animals. Wilcoxon signed-rank test. c) Example recording during REMS-R, with grey rectangle indicating portion expanded on the right. Mean transitional dynamics calculated as in Fig. 2a. d) Example recording at ZT1 and at ZT5, corresponding to early and late moments during the REMS-R. e) e1) Quantification of mean (inter-REMS) intervals across a 6 h-REMS-R, with mean values shown on the right; e2) Same for LC activity levels in the 5 s before REMS entries. Grey lines connect paired data per animal. Paired t-tests for e1 and e2. f) Example recording for a REMS-R combined with Sham (LED-off) or optogenetic stimulation of LC at 1 Hz. Quantification of REMS entries across the stimulation period. Note the higher REMS transition frequency before optogenetic stimulation compared to baseline sleep (Fig. 2 b, c). Box-and-whisker quantification of mean data for the entire REMS-R. Wilcoxon signed-rank test. See **Supplementary Table 1** for detailed statistical information.

We examined the timing of REMS entries with respect to fiber photometric measures of LC activity during REMS-R in a separate group of 8 animals. The mean duration of REMS bouts during the REMS-R was 8.9 ± 2.4 s (n = 8), similar to the recently reported values of another automated REMS-R method^43^. During REMS-R, every REMS entry remained invariably preceded by a decline of LC activity (**Fig. 3c**). Furthermore, although REMS-R shortened the time between consecutive REMS entries, these remained locked to moments of low LC activity levels (**Fig. 3d**). Starting from the 4^th^ h of the REMS-R, the mean interval remained comparatively stable and settled at 48.2 ± 6.3 s at ZT6 (**Fig. 3e1**). To examine the importance of low LC activity levels for REMS entries, we calculated corresponding values for the last 5 s before every REMS entry across the full time of REMS-R. We found these levels to remain stably low and constant during the entire 6 h of REMS-R (**Fig. 3e2**). This strongly suggests that the decline of LC activity to a minimal level remained obligatory for a REMS entry to become possible, in spite of a substantial increase in the pressure to enter REMS. To directly test whether LC activity could suppress REMS entries even when the pressure for REMS was elevated, we combined optogenetic LC stimulation in ChR2-expressing DBH-Cre animals with REMS-R (n = 6, **Fig. 3f**). LC was stimulated at 1 Hz for 20 min/h, known to suppress baseline REMS (see **Fig. 2b**). This LC stimulation also suppressed REMS entries during the entirety of the REMS-R, with sham (LED-off) conditions showing no effect. This indicates that moderate levels of LC activity (1 Hz) can override a high REMS pressure to suppress REMS entries.

The combined evidence of a novel REMS-R method together with LC activity monitoring or optogenetic testing supports an obligatory role of LC activity decline in ruling REMS entries, even under conditions of high REMS pressure. The infraslow time course of LC’s natural alternation between high and low activity levels thus presents a previously unrecognized time interval that gatekeeps sleep’s organization in NREMS-REMS cycles.

### LC activity fluctuations gatekeep the natural NREMS-REMS cycle duration

The mechanisms regulating natural cycle length remain poorly understood, although homeostatic mechanisms have been implied^42^. Here, we asked whether the LC-associated infraslow activity fluctuations were relevant for natural cycle length in undisturbed mouse sleep^44^. In mouse, times spent in NREMS between REMS vary in length from < 1 min to > 30 min^44^. To probe whether LC inhibition can modify cycle length, we therefore aimed to identify the set of cycles with relatively little variability in duration. Following previous investigations on the interrelationship between NREMS and REMS duration in mouse sleep^44,45^, we analyzed how the time spent in NREMS depended on prior REMS time in the light phase (n = 17). This showed that long REMS bouts (> 120 s) were most consistently followed by long NREMS bouts, whereas shorter (< 100 s) REMS bouts were followed by variable times spent in NREMS (**Fig. 4a, b**). In undisturbedly sleeping Jaws-expressing DBH-Cre mice (n = 7), we hence measured the duration of each REMS bout in real-time, and, once it lasted ≥ 120 s, we triggered optogenetic inhibition of LC in a double closed-loop experimental design. The onset of this inhibition was set ‘early’ (within 50-100 s after NREMS onset) or ‘late’ (after NREMS time lasting twice the duration of the preceding REMS bouts ± 10 s) time points in NREMS, which allowed to probe LC’s role at different moments in the NREMS-REMS cycle. Sham (LED-off) sessions took place interleaved with test (LED-on) sessions (see Methods). When LC inhibition started early in NREMS, we observed no significant change in the frequency of REMS entries or in times spent in REMS (**Fig. 4c**). In contrast, when LC inhibition started late, repeated REMS entries took place almost twice as frequently compared to Sham stimulation (**Fig. 4d**). We confirmed the efficiency of inhibition by histological examination of the correct optic fiber location (**Extended Data Fig. 5a**). To obtain an additional independent measure for this efficiency, we examined how the observed increases in REMS transitions correlated with changes in sleep spindle densities (**Extended Data Fig. 5b**)^17^. This yielded a positive correlation, which supported the idea that a more efficient optogenetic LC inhibition allowed for a stronger augmentation of REMS entries. As a final control, we demonstrated that mCherry-only-expressing controls (n = 7) did not change the frequency of REMS entries upon light exposure (**Extended Data Fig. 5c**). Natural LC activity fluctuations are thus critical for the normal NREMS-REMS-cycle, in part because they appeared to prevent precocious REMS entries late in the cycle and prolonged the time spent in NREMS. In contrast, LC activity appears to play a minor role at relatively early stages of the cycle, when a long REMS bout was just completed and for which a refractory period has been proposed^44^.

**Figure 4.**
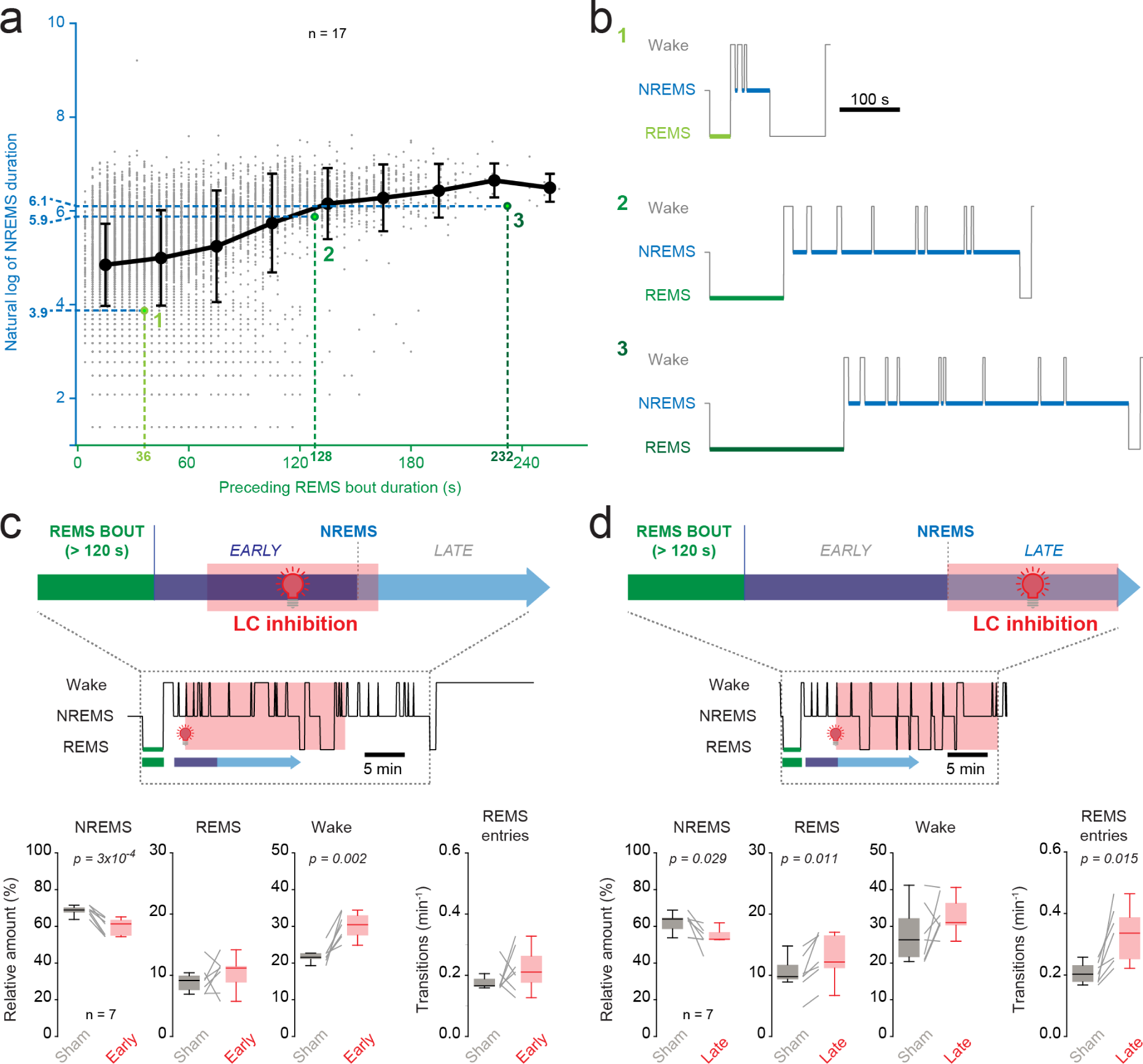
LC activity declines were necessary for REMS entries late but not early in the natural NREMS-REMS cycle. a) Scatter plot of NREMS times (in units of natural logarithm) *vs* preceding REMS bout durations for undisturbed sleep of n = 17 animals in the light phase, as done in ref^44^. Datapoints marked by 1, 2 or 3 mark examples shown in panel b. b) Representative examples for datapoints highlighted with numbers in panel a. c) Top, schematic of the LC optogenetic inhibition protocol starting early in the NREMS bout once a REMS episode lasting > 120 s occurred. Example hypnogram is presented below. Box- and-whisker plots quantify times spent in wake, NREMS and REMS, and the frequency of REMS entries for early light inhibition (n = 7). Wilcoxon signed-rank or paired t-test. d) As e, for LC inhibition starting late in the NREMS bout (n = 7). Wilcoxon signed-rank or paired t-tests. See **Supplementary Table 1** for detailed statistical information.

### A stimulus-enriched wake experience increased LC activity fluctuations in NREMS and disrupted its architecture

Given the implication of LC activity fluctuations for the NREMS-REMS cycle, modifications in the strength of these fluctuations could powerfully disrupt sleep. A recent study found that acute stress during wakefulness can severely perturb LC activity in subsequent NREMS^19^, but stress effects and alterations in sleep-wake behavior remain to be dissociated. Here, we subjected jGCaMP8s-expressing DBH-Cre mice to two different 4 h-long sleep deprivation (SD) procedures from ZT0-ZT4. One SD was achieved by the ‘gentle handling’ technique that prevented animals from falling sleep while minimally disturbing their spontaneous activity patterns in the cage (**Fig. 5a**). The stimulus-enriched SD (SSD) was instead paired with auditory and somatosensory stimuli, to which the animals were not previously habituated and exposed to in an unpredictable manner, thus promoting LC activation (**Fig. 5b**)^1^. Both SD and the SSD were carried out at least 10 d apart in groups of 4 animals. Blood sampling immediately after the SSD in a separate group of animals revealed increases in corticosterone levels compared to baseline (**Extended Data Fig. 6**), indicating elevation of stress levels in addition to the sensory exposure, which strongly implicates LC^1,2^. The SD produced by gentle handling instead does not increase corticosterone levels^46^.

**Figure 5.**
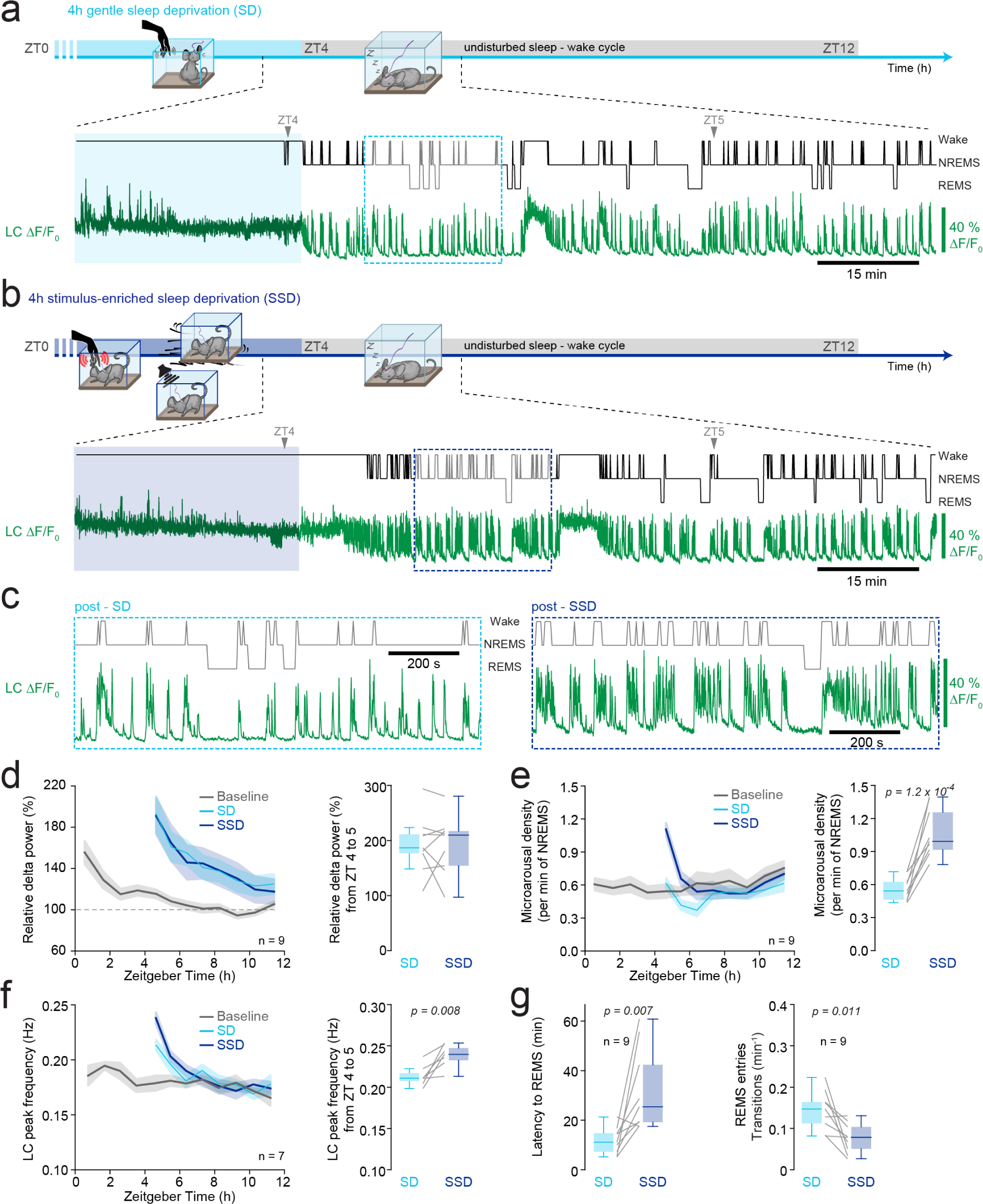
A stimulus-enriched but not a gentle sleep deprivation disrupted subsequent sleep and enhanced LC activity. a) Schematic of experimental timeline for gentle sleep deprivation (SD), with example hypnograms and corresponding LC fluorescence from the the end of the SD to 4 h in post-SD. Dotted rectangle indicates expanded portion in c. b) As a, for the stimulus-enriched sleep deprivation (SSD). Dotted rectangle indicates expanded portion in c. c) Expanded portions of traces in a, b. d) Mean dynamics of EEG δ power from ZT0-ZT12 for n = 9 animals subjected to baseline recordings, SD and SSD. Box-and-whisker quantification for data from ZT4-ZT5 on the right, corresponding to the first h after SD or SSD, with grey lines showing paired data per animal. Paired t-test. e) As c, for the density of MAs (n = 9). Paired t-test. f) As c, for LC peak frequency (n = 7). Paired t-test. g) Box-and-whisker quantification for REMS onset latency, quantified from the time of NREMS onset, and the transitions frequency of REMS entries, for the first h after SD or SSD (ZT4-ZT5, n = 9). Paired t-tests. See **Supplementary Table 1** for detailed statistical information.

As evident from the two representative recordings of sleep-wake behavior immediately after SD or SSD of the same animal (**Fig. 5a-c**), LC continued fluctuating between activity transients and troughs during NREMS, yet with the former ones distinctly strengthened. Both SD and SSD resulted in comparable increases in low-frequency power of the EEG δ (1.5 – 4 Hz) band, indicating that the homeostatic pressure to sleep had become elevated similarly in response to the 4-h wake time (**Fig. 5d**). Remarkably, the density of MAs changed in opposite directions for SSD compared to SD. SD decreased MA frequency in subsequent NREMS, consistent with homeostatic sleep consolidation^32^. The SSD instead resulted in a marked increase in MA density and hence fragmented NREMS (**Fig. 5e**). Furthermore, LC activity, measured in peak frequency (see Methods), was higher in the first h of SSD compared to SD (**Fig. 5f**). Finally, the latency to REMS onset was increased in SSD compared to SD and the number of REMS entries decreased within ZT4-ZT5 (**Fig. 5g**). In comparison to an extended period of wakefulness, a stimulus-enriched wake experience combined with stress transiently increased LC activity, disrupted sleep architecture for a post-SD/-SSD period of 1-2 h, and antagonized the homeostatic effects of sleep consolidation.

### Inhibiting LC activity antagonized sleep disruptions induced by a stimulus-enriched wake experience

We examined the role of LC activity fluctuations in the SSD-induced sleep disruptions. First, we determined the spectral profiles associated with high and low LC activity levels, as done for baseline recordings (see **Fig. 1h, i**). LC activity transients remained associated with characteristic spectral changes in the EEG for MA-associated events, however, increases in δ (1.5 – 4 Hz) power for non-MA associated events were suppressed (**Fig. 6a**) in comparison to the SD condition (n = 7, paired t-test, p = 0.038, effect size D = −1.42). All REMS entries also remained invariably associated with LC activity declines (n = 7, **Fig. 6b**). Second, to causally test the consequences of elevated LC activity, we subjected 9 Jaws-expressing DBH-Cre animals to two SSDs spaced at least 10 d apart, of which one was followed by 1 or 2 h of Jaws-mediated bilateral inhibition of LC, the other one by LED-off (sham) exposure, in a counterbalanced manner. LC inhibition attenuated the fragmentation of NREMS and shortened REMS onset latency (**Fig. 6c**). LC’s activity fluctuations in NREMS thus depend on the type of wake experience and their modification can increase spontaneous arousability and delay REMS. These data suggest that abnormal LC activity fluctuations can be causally involved in two important hallmarks of deteriorated sleep quality.

**Figure 6.**
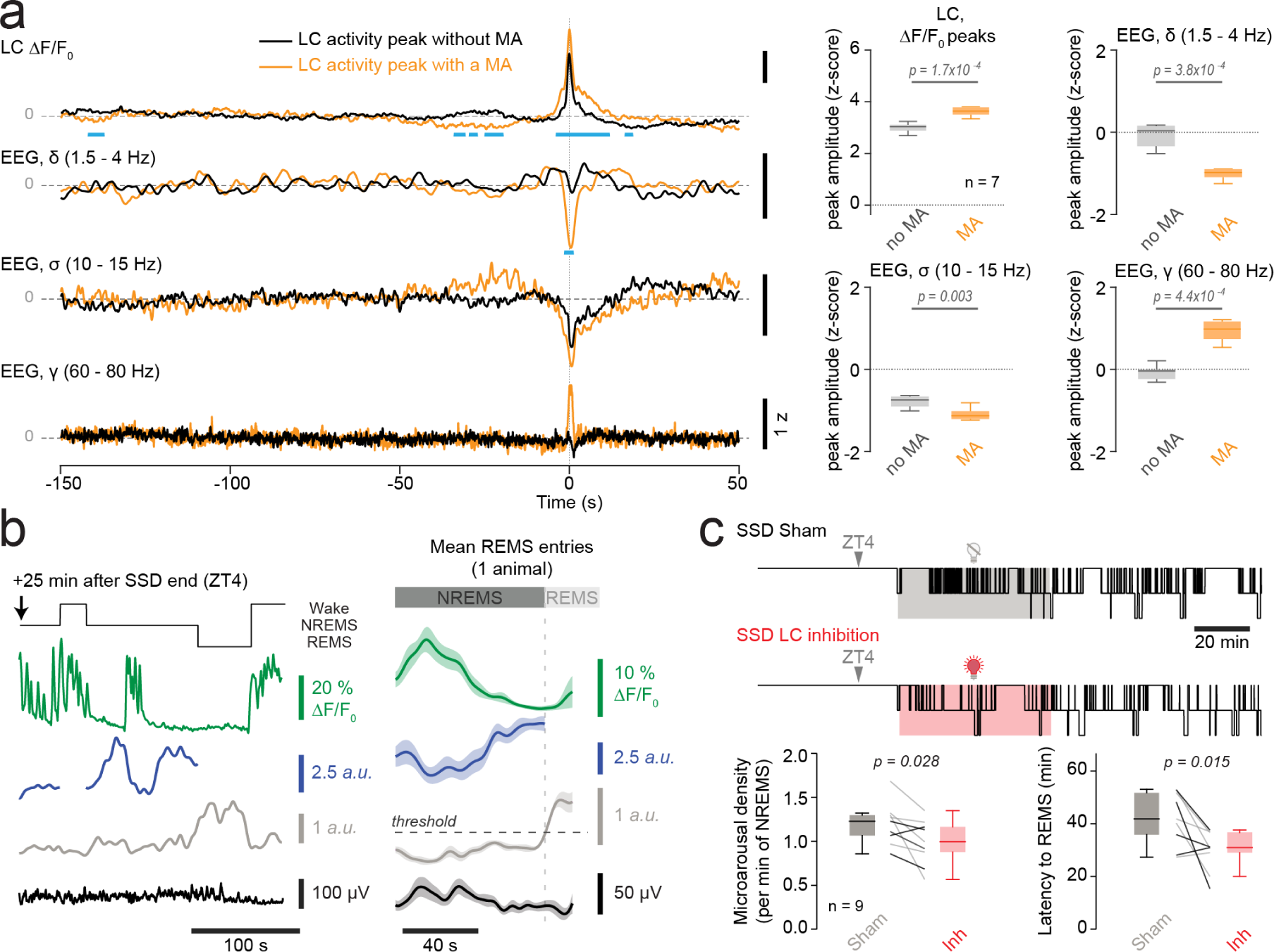
Acute LC inhibition suppressed sleep-disruptive effects of a stimulus-enriched wakefulness. a) Spectral analysis of EEG signals corresponding to LC activity peaks, as done in Fig. 1 h, I for the first 2 h after the end of the sensory stimulus-enriched sleep deprivation (SSD). LC activity peaks that are classified based on whether they occurred without a MA (black) or with a MA (orange). Traces denote means across n = 7 animals. Blue bars denote significance p values as calculated by false discovery rates. Quantification of mean values from traces shown in a for peak values for the time interval 0 – 1 s. All p values were calculated from paired *t*-tests. For power analyses, the Bonferroni-corrected p = 0.017. b) Example recording of a REMS entry after SSD. Mean transitional dynamics calculated as in Fig. 2a. c) Example hypnogram after SSD with or without Jaws-mediated optogenetic inhibition of LC. Box-and-whisker quantification of MA density and REMS onset latency after SSD with sham (LED-off) or light stimulation (n = 9). Grey and black lines connect data from animals exposed to 1-h or 2-h light-induced LC inhibition post-SSD. Paired *t*-test. See **Supplementary Table 1** for detailed statistical information.

## Discussion

We identify the noradrenergic LC as a gatekeeper for the organization of mammalian sleep into NREMS-REMS cycles. The underlying principle of action is LC’s partitioning of NREMS into two distinct brain and autonomic states, of which one was arousal-promoting, the other one permissive for REMS (**Figure 7**). To date, we know much about the neural circuits underlying NREMS and REMS individually, and both play important roles for sleep’s restorative and memory-promoting functions^47–49^. In comparison, the circuit mechanisms that rule the alternation between NREMS and REMS have remained open, despite pioneering modeling and experimental work^11,24,25,41,42,50^. The identification of the LC as a gatekeeper for the NREMS-REMS cycle is conceptually novel because it points to an upstream oscillator-like mechanism that sets the time windows for the operation of NREM-or REMS-regulating circuits. We also demonstrate that disrupting this gatekeeper fragmented sleep and disrupted the NREMS-REMS cycle, and that it could override the consolidating effects of NREMS homeostasis.

**Figure 7.**
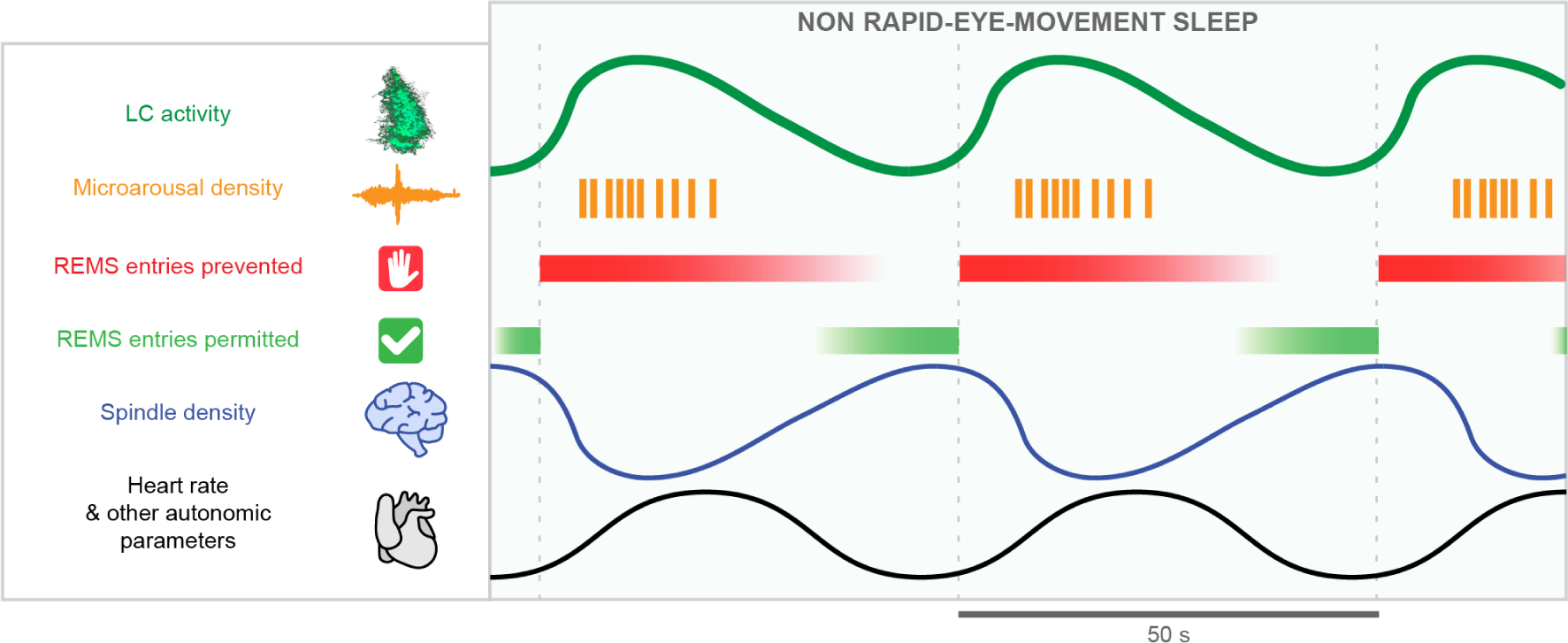
Scheme summarizing the functional partitioning of NREMS and its relevance for the NREMS-REMS cycle From top to bottom, the time course of physiological parameters relevant for the partitioning of NREMS into arousability-promoting/REMS-supressing and REMS-promoting periods is illustrated. Two full cycles of this functional partitioning are shown, indicated by vertical dashed lines. Green thick lines: LC neuronal population Ca^2+^ activity; Orange: Frequency of MAs (see also ref ^21^); Red and Green Stop and Go signals: REMS-suppressing and REMS-permissive periods; Blue line: Brain spectral fluctuations in sigma power (see also refs ^17,22^); Black, Heart rate fluctuations (see also refs ^17,22^). Other autonomic parameters include pupil size fluctuations as demonstrated in ref^72^.

The LC was included as a REMS-off (that is, REMS-suppressing) brain area in the first theoretical models for the NREMS-REMS cycle^24,25^, which has been supported experimentally^17–19^. Through timing optogenetic interference to LC’s natural activity fluctuations, we show here that LC plays a dual role, suppressing REMS on the one hand, and setting the moments when REMS entries are permitted on the other hand^25,42^. LC’s permissive role adds an infraslow periodicity to the architectural control of the NREMS-REMS cycle, defining recurring moments during which a REMS entry is possible^42^. A possible advantage of LC’s permissive role is an adaptability of NREM-REMS cycle duration to circadian and homeostatic regulation, developmental state, metabolic and emotional conditions, all of which shape the mammalian NREMS-REMS cycle across the lifespan^51^. Our study opens the way to explore these possibilities, for example by investigating how infraslow LC activity fluctuations interplay with REMS-promoting areas in hypothalamic, brainstem, or mesolimbic circuits.

This study further shows that unique brain and autonomic states associated with LC activity fluctuations are core for the functional partitioning of NREMS. Following earlier work^16–18,22^, we find that LC activity transients acted in an arousal-promoting manner for the autonomic nervous system (through accelerating heart rate) and for thalamic sleep spindle-generating circuits (through suppressing sleep spindles)^17^. This is in line with an LC promoting arousal during NREMS. Unexpectedly, we identify a cortical ‘sleep-like’ EEG or LFP activity that coincided with thalamic and cardiac activation, suggesting that LC-linked arousal promotion involved subcortical but not cortical areas. We show this subcortical arousal here for the sensory thalamus, but we speculate that other areas strongly innervated by LC, such as midbrain and cerebellum^52^, as well as other REMS-regulating areas^7^, might also be recruited, which could underlie the prevention of entries into REMS. In support of a subcortically delimited action of LC, a functional imaging study in anesthetized rats indicated that low-frequency optogenetic LC stimulation increased BOLD signals in the thalamus but not in the cortex^53^. Furthermore, a consistent thalamic activation yet a variable cortical involvement was identified via intracranial recordings for MAs in human NREMS^54^. Our analysis thus advances a circuit foundation for a heterogeneity of aroused brain states in which wake-and sleep-related brain activity can coexist. This will be relevant in dissecting the temporal sequence of arousal processes to full wakefulness^55^ and its possible dysfunction in disorders of arousals^56^.

We further propose that the spectral and autonomic profile of LC activity is valuable in identifying the diversity of arousals in the human sleep EEG. There are established arousal scoring guidelines for human sleep^57^, nevertheless, there remain EEG signatures for which the relationship to arousal remains debated^23^. One example is a singular, large low-frequency EEG wave that is prominent in human NREMS stage 2, referred to as ‘K-complex’^58^. K-complexes are associated with small but consistent increases in heart rate^59,60^, yet they have been variably described as sleep-protecting or arousal-related EEG hallmarks^58^. The data we present now indicate that increases in low-frequency activity in the EEG can be a forebrain manifestation of activated brainstem arousal circuits, which is in line with suggestions on the origins of K-complexes based on EEG source analyses^61^. We suggest that some K-complexes in human sleep should be examined with respect to a possible implication of LC activity in human NREMS. A recent study using pupillometry as a proxy for LC activity in human sleep demonstrates that such progress is in reach^60^.

Possible neural mechanisms by which LC activity activates the thalamus while promoting low-frequency activity in the cortex can be derived from available knowledge on NA’s actions within thalamocortical circuits. Single LC unit recordings from rats in NREMS showed that individual spikes are phase-locked to cortical slow (low-frequency) waves^13^, while only high-frequency LC stimulation activates the cortex^53^. Therefore, the NA release generated by minor neuronal discharge typical for NREMS appears insufficient to disrupt the bistability of cortical up and down states, while it depolarizes thalamocortical and thalamic reticular neurons and suppresses burst firing^17^. An excitatory drive to the cortex that would result from enhanced thalamic output could prompt switches between up and down states of cortical circuits^62^, which would manifest as a transient increase in low-frequency power. Once LC activity transients are longer or larger, as is for example the case after SSD, its impact on cortex could be elevated and low-frequency power become suppressed. Other mechanisms involving target-specific synaptic release of NA and peptide messengers need to be further tested^1^.

The important role of LC activity fluctuations for the NREMS-REMS cycle could be demonstrated by asking how frequently entries from NREMS to REMS could be maximally permitted. For this, we developed a specific REMS-R procedure as a next-level refinement of automated approaches for REMS restriction^43^. This allowed to directly visualize that LC activity fluctuations persisted even during high REMS pressure and that every REMS entry coincided with an invariable, low, LC activity. Minimal, non-reducible time intervals were previously reported in REMS-restriction protocols of similar duration^63,64^, yet we provide here a first mechanistic basis that raises novel questions about the role of this minimal duration. We cannot currently exclude that mechanisms besides LC play a role, but we note that the noradrenergic LC is implied in neuronal, glial, and vascular signaling mechanisms in the brain^7^, all of which might take time to complete in NREMS before REMS can start. We further note that REMS entries occurred in an almost oscillatory-like manner in the last hours of the REMS-R, during which REMS pressure became excessively high. In reptile sleep, alternations between NREM-and REMS-like states recur periodically on the infraslow time scale^51,65^. We think that the infraslow time scale could be an evolutionarily conserved time interval for the NREMS-REMS cycle over which a gatekeeping mechanism evolved later to adapt the NREMS-REMS cycle to a variety of physiological needs.

An aversive or stressful experience in humans may disrupt sleep, for example by fragmenting sleep and by suppressing REMS^66,67^. Several recent studies have identified the LC to be involved in sleep disruptions preceded by acute stressful experiences^19,68^, illustrating the growing recognition of neural circuits involving monoamine release as a source for sleep disturbances. As LC controls both, spontaneous arousability and REMS gatekeeping, alterations in its activity could have significant consequences. Indeed, we found that a transient negative wake experience was sufficient to disrupt the balance of LC activity fluctuations towards sustained high levels of activity. The resulting sleep fragmentation and delay in REMS onset could be attenuated by antagonizing these elevated levels of LC activity. This establishes the LC as an area that makes sleep vulnerable to recent, even transient, and moderately adverse wake experiences.

The study offers a neural mechanism relevant for the alternation of mammalian sleep between states of NREMS and REMS. Beyond this fundamental insight, our work has opened a translational research line that will strengthen the connection between rodent and human sleep because it proposes novel testable biomarkers for LC activity in human sleep. Furthermore, elucidating the origins of LC’s gatekeeping role could shine light on certain NREMS functions that inevitably require time or coordinated to complete before a transition to REMS can be made. The susceptibility of sleep-active LC to the preceding wake experience further brings novel opportunities to elucidate sleep disruptions that could be impacted by the diversity of our daily experiences.

## Acknowledgements

The authors acknowledge fruitful discussions with Paul Franken, Stephany Fulda, Caroline Lustenberger, Markus Schmidt, Francesca Siclari, Aurélie Stephan, Mehdi Tafti and Eus van Someren. We thank Dr. Simone Astori and Olivia Zanoletti from the Carmen Sandi lab (EPFL, Lausanne) for carrying out the corticosterone measurements and Aurélie Guillaume-Gentil for assistance with viral injections. We recognize Dr. Sandro Lecci for providing the first datasets relevant for this study. Many thanks to Drs. Feng and Li for generously providing us with the first aliquots of the GRAB_NE_ sensor. The constructive comments of Dr. Julie Buron, Lila Banterle and Paola Milanese on the manuscript are very appreciated. We are grateful to the animal caretakers, especially Michelle Blom and Titouan Tromme, and to Dr. Laure Sériot for expert veterinary advice. This work was supported by the Swiss National Science Foundation (Grant no. 184759 to AL), a FBM UNIL Doctoral Fellowship to AOF, and Etat de Vaud.

## Materials and Methods

### ANIMALS AND HUSBANDRY

The study used two mouse lines - the C57BL/6J line and the B6.FVB(Cg)-Tg(Dbh-cre)KH212Gsat/Mmucd (MMRRC Stock#036778-UCD) line, referred to here as DBH-Cre line, that was originally provided to us by David McCormick and Paul Steffan, University of Oregon. Mice were bred on a C57BL/6J background and housed in a humidity-and temperature-controlled animal house with a 12 h / 12 h light-dark cycle (lights on at 9 am, corresponding to ZT0). Food and water were available *ad libitum* throughout all experiments. For viral injections, 3-to 10-week-old mice of either sex were transferred to a housing cabinet in a P2 safety level room, where they stayed from 1 d before to 3 d after the viral injection. They were then transferred to the sleep-wake recording room and left to recover for ≥ 1 week before undergoing the implantation surgery, after which they were single-housed in standard-sized cages. The grids on top of the cage were removed and replaced by 30 cm-high Plexiglass walls. Fresh food was regularly placed on the litter and the water bottle inserted through a hole in the cage wall. Objects (tissues, paper rolls, ping-pong balls) were given for enrichment.

In total, 10 DBH-Cre mice were used for two 12-h baseline sleep recordings in the light phase in combination with fiber photometry under undisturbed conditions. One animal was excluded because the dynamic range of the fluorescent signal (ΔF/F_0_ < 15%, see below). Additionally, two of these 2 DBH-Cre animals were used for combined fiber photometric monitoring of LC activity and NE levels, these were included in the analysis of LC fluorescence across wake, NREMS, REMS. For combined LC optogenetic inhibition with fiber photometric monitoring of NE release, 2 DBH-Cre animals were used. Of the 10 DBH-Cre animals included for fiber photometric monitoring of LC activity, 8 were used for 1-2 sessions of REMS-R in combination with fiber photometric monitoring of LC activity, and 9 underwent 1-2 sessions of SD and one session of SSD in combination with fiber photometric monitoring of LC activity. Animals in which the dynamic range of fluorescence fell < 15% during repeated fiber photometric sessions were no longer included in the analysis of LC activity. 9 DBH-Cre mice for optogenetic stimulation and 9 DBH-Cre mice for optogenetic inhibition during different phases of the infraslow fluctuation in σ power were already used in an earlier study and re-analyzed here^17^. Seven DBH-Cre mice were used for optogenetic inhibition early or late in the NREMS-REMS cycle, and an additional 7 were used for control viral injections. For the analysis of the statistics of the NREMS-REMS cycle, baseline or sham sleep recordings from these 14 animals in the light phase, plus additionally 3 animals otherwise not included in these experiments, were included. For the validation of the REMS-R procedure alone, 9 C57BL/6J animals were used. For REMS-R in combination with optogenetic stimulation, 6 DBH-Cre animals were used. 9 animals were used for SSD in combination with optogenetic LC inhibition. A separate group of 8 animals were used for quantification of corticosterone levels at ZT4 in baseline and after SSD. All experiments were conducted in accordance with the Swiss National Institutional Guidelines on Animal Experimentation and were approved by the Swiss Cantonal Veterinary Office Committee for Animal Experimentation.

### VIRAL VECTORS AND INJECTIONS

General and local anesthesia, analgesia, temperature control, stereotaxic fixation and drilling of craniotomies for injections of viral vectors were as described^17^. For fiber photometric monitoring, DBH-Cre mice were unilaterally injected over the right LC (L 1.05; AP −5.4; DV −3.2) with AAV5-hSyn1-dlox-jGCaMP8s-dlox-WPRE-SV40p(A) (titer 5.8E×10^12^ vg/ml, 300nL, unilateral, Viral Vector Facility (VVF) Zürich) at an injection rate of 50-100 nL/min using a thin glass pipette (5-000-1001-X, Drummond Scientific) pulled on a vertical puller (Narishige PP-830), initially filled with mineral oil, and backfilled with the virus-containing solution just prior to injection. For transfection of the LC for optogenetics, viral vectors injected were ssAAV5/2-hEF1α-dlox-hChR2(H134R)_mCherry(rev)-dlox-WPRE-hGHp(A) (9.1×10^12^ vg/ml, 0.8 – 1 μL, unilateral, VVF Zürich) or ssAAV-5/2-hSyn1-dlox-Jaws_KGC_EGFP_ERES(rev)-dlox-WPRE-bGHp(A)-SV40p(A) (6.4×10^12^ vg/ml, 300 nL bilateral, VVF Zürich). The control virus for Jaws experiments was ssAAV-5/2-EF1α-dlox-mCherry(rev)-dlox-WPRE-bGHp(A) (7.3×10^12^ vg/ml, 300 nL bilateral, VVF Zürich). For monitoring of free NA levels, ssAAV9/2-hSyn1-GRAB_NE1h-WPRE-hGHp(A) (7.2×10^12^ vg/ml; VVF Zürich) was injected unilaterally in primary somatosensory thalamus (500 nL; L 2.0; AP −1.6; D −3.0), as described before ^17^. Two animals each were dually injected for combined expression of jGCaMP8s and GRAB_NE1h_ or Jaws and GRAB_NE1h_ in the right hemisphere. After the injections, the incision was sutured and disinfected, and the animal was supervised and given paracetamol at 2 mg / ml for the next 4 days.

### SURGERIES

#### Surgeries for sleep recordings

Surgeries for implantation of electrodes for sleep-wake monitoring were done at least 1 week after the viral injection. Following anesthesia induction (5% Isoflurane, animals were gas-anesthetized (Isoflurane 1.5 – 2.5%) in a mixture of O_2_ and N_2_O. After analgesia (Carprofen 5mg/kg i.p.) and disinfection, animals were fixed in a Kopf stereotax and injected into the scalp with a mix of lidocaine (6 mg/kg)/bupivacaine (2.5 mg/kg). To expose the skull, the skin was incised, and the bone scratched with a fresh scalpel blade to improve adhesion of the head implant. Then, we drilled small craniotomies (0.3 – 0.5 mm) over left frontal and parietal bones and positioned two conventional gold-coated copper-wire electrodes in contact with the dura mater for EEG recordings. In case of unilateral LC injections, EEG electrodes were implanted contralaterally. On the ipsilateral (right) side, high-impedance tungsten LFP microelectrodes (10–12 MΩ, 75-μm shaft diameter, FHC) were implanted in the primary somatosensory cortex (L +3.0; AP −0.7; D −0.85) and the hippocampus (L +2.0; AP −2.46; D −1.2 to −1.3). Additionally, as a neutral reference, a silver wire (Harvard Apparatus) was inserted into the occipital bone over the cerebellum and two gold pellets were inserted into the neck muscles for EMG recordings. All electrodes were fixed using Loctite Schnellkleber 401 glue and soldered to a multisite connector (Barrettes Connectors 1.27 mm, male connectors, Conrad). Animals were single housed after surgeries.

#### Surgeries for fiber photometry recordings

For fiber photometry, fiber implantation occurred together with the surgical implantations for sleep recordings. Shortly, an optic fiber stub coupled to a cannula (Doric Lenses, MFC_400/430-0.66_3.5mm_ZF1.25(G)_FLT) was implanted over the right LC (L +0.9; AP −5.4; D −2.7) at an insertion speed of 1 mm/min using a Kopf stereotaxic holder, for 1 animal the fiber was implanted with an angle of 20 degrees leading to a different set of coordinates (L +1.84; AP −5.4; D −2.9). For dual fiber photometry, this procedure was repeated over the primary somatosensory thalamus (L +1.8; AP −1.7; D-2.5), over which the fiber stub was vertically inserted. Fibers were glued to the skull and were part of the dental cement structure holding the entire implant into place.

#### Surgeries for optogenetics

For optogenetic stimulation of noradrenergic LC neurons, 4 mm-long optic fibers were purchased pre-pared (MFC_200/250-0.66_4mm_ZF1.25(G)_FLT) or fabricated from a multimode fiber (225 μm outer diameter, Thorlabs, BFL37-2000/FT200EMT) using the custom-made procedure described in ref^17^. For optogenetic inhibition, 5 mm-long optic fibers attached to an optic canula were used (CFMLC12L05, Thorlabs). The implantation coordinates for stimulation were (L +0.9; AP −5.4; D −2.5 unilaterally in the right LC), while for inhibition two fibers were implanted bilaterally using a 20° angle from the vertical with coordinates (L +/-1.84; AP −5.4; D −2.5) from bregma. The emitted light intensity at the fiber tip was evaluated prior to implantation for every fiber.

### RECORDING PROCEDURES AND PROTOCOLS

#### Sleep recordings

Sleep recordings always took place within the animals’ home cage. Once recovered from surgery for > 1 week, animals were habituated to cabling to an ultrathin Intan SPI cable and adaptor board via a custom-made support system (Homemade adapters containing an Omnetics A79022-001 connector linked to a female Barrettes Connector from Conrad). These served as intermediate between the head implant of the animal and the headstage (RHD2132 amplifier board) that was connected to the Intan USB interface Acquisition Board (RHD2000). All signals (EEG frontal and parietal, left and right EMG, S1 and CA1 LFPs) were acquired in unipolar mode at 1 kHz.

All recordings were done in the 12-h light phase (ZT0-ZT12). For closed-loop optogenetic stimulation and inhibition sessions timed to the infraslow σ power fluctuations in NREMS, 2 sessions were done per condition (Rising or declining σ power, Sham LED-off or Light stimulation). In experiments using optogenetic stimulation or inhibition according to the infraslow phase, sessions were spaced by one day and data meaned between sessions. For optogenetic inhibition timed to the early or the late phase of the NREM-REMS cycle, 10 to 20 sessions per animal were recorded over 3 weeks’ time. For the two REMS-R sessions per animal, at least 2 interleaved days were given. Per animal, there were 1-2 SDs that were kept at least 7 days apart. The SSD was spaced apart from the SD sessions by at least 7 days, and 10 days were given for two SSDs combined with LC optogenetic inhibition or sham (LED-off) exposure. For details of the light exposure, see ‘Optogenetic protocols’.

#### Fiber photometry recordings

Fiber photometry recording sessions took place at least one week after surgery and >1 day apart. Two signal generators modulated the blue (465 nm) and the violet (405 nm) LED of the Doric Fluorescence MiniCube LEDs (ilFMC4-G2_IE(400-410)_E(460-490)_F(500-550)_S) using sinusoidal signals at 319 and 211 Hz, respectively. The combined modulated light was then transmitted through a 400-μm-thick fiberoptic patchcord (MFP_400/430/1100-0.57_1m_FMC-ZF1.25_LAF, Doric Lenses) to the implanted optic canula on the head of the animal. A photodetector integrated into the MiniCube turned the emitted light from the fluorescent sensors into a current signal that was fed into an analog signal of the Intan RHD2132 amplifier board as described previously^17^. The measured power at the tip of the optic fiber was, for both wavelengths, 20 – 30 µW ensuring minimal bleaching throughout the 12 h of recordings.

#### Optogenetics protocols

For optogenetic interference at different phases of the infraslow σ power fluctuations, protocols were applied as described^17^. The optogenetic manipulations took place every first 20 min/h from ZT1-ZT9. Stimulation sessions (with LED on) took place in exchange with sham session (LED off). Entries into NREMS were detected online through setting a threshold for the low-frequency (1 – 4 Hz) over θ (6 – 10 Hz) ratio for the differential frontal-parietal EEG signal and the decline in the absolute EMG amplitude. Occasional brief interruptions in LED stimulation during NREMS are due to a trespassing of that ratio, either caused by muscle twitches or spectral fluctuations. Once in NREMS, σ power fluctuations were monitored online and increasing or decreasing slopes detected through a machine-learning procedure^17^. For LC stimulation, we used 1 Hz pulses of 10 ms applied to a PlexBright Plexon BlueLED unit emitting 465 nm blue light at 2.8-3.2 mW intensity. For LC inhibition, we used continuous stimulation of an orange LED module (620 nm, 1.55 – 1.7 mW at the tip).

For optogenetic interference at different time points in the NREMS-REMS cycle, we used a closed-loop algorithm to identify REM and NREMS based on the low-frequency (1 – 4 Hz) over θ (6 – 10 Hz) ratio for the differential frontal-parietal EEG signal and the EMG activity based on 4-s windows with a 3 s overlap (i.e. updated every s), as previously described^21^. Upon online identification of long REMS bouts (>120 s), 20-min continuous optogenetic inhibition started randomly after 50 – 100 s of NREMS (*early inhibition*), or after twice the time of the identified REM bout (at least 240 s or 2 times the duration of the preceding REMS bout) with a random 10-s jitter (*late inhibition*). The time points chosen to start the early or the late inhibition were based on a previous study analyzing the probabilities for REMS entries after a long REMS bout^44^. Several sessions were recorded so that we collected around 10 episodes of long REM bouts per condition per animal.

For optogenetic inhibition after SSD (see below for procedure), LC was continuously inhibited for 1 or 2 h after the end of the SSD using continuous illumination. The stimulation started once a 10-s long NREMS was automatically detected, using a closed-loop algorithm based on the low-frequency (1 – 4 Hz) over θ (6 – 10 Hz) ratio for the differential frontal-parietal EEG signal and the EMG activity. This resulted in a mean of 36.3 ± 5.1 min and 34.5 ± 4.5 min of sleep/h of light stimulation for sham and test conditions, respectively (n = 9). One additional animal was *post hoc* excluded from analysis because it slept only ∼5 min during the 2-h post SSD light exposure, which provided insufficient data for analysis. Per animal, one SSD followed by LC inhibition and one SSD followed by Sham inhibition was carried out in a counterbalanced design.

### BEHAVIORAL MANIPULATIONS

#### REMS restriction (REMS-R)

We used small vibrating motors (DC 3–4.2 V Button Type Vibration Motor, diameter 11 mm, thickness 3 mm) that we fixed using double-sided tape to the end of the recording cables, close to the animals’ heads. The motors were driven to vibrate through a closed-loop system using a Raspberry Pi^21^. Once the θ (6 – 10 Hz) / δ (1 – 4 Hz) ratio of the CA1 LFP signal passed a threshold 1.5 for > 2 s and EMG activity declined below a threshold of 0.35 that was based on the min/max normalization of the median log10 EMG activity/s, the motors vibrated for 2 s and woke up the animal, as monitored by polysomnography. A control session for protocol standardization was included in which the motor of an animal was activated based on the closed-loop of the neighbor animal (yoked controls). The animals included only for the validation for the REMS-R procedure were separate from the ones in which LC activity was monitored or optogenetically altered during the REMS-R in terms of analysis. The REMS-R was carried out from ZT0-ZT6 for all experiments except when REMS-R was combined with LC optogenetic stimulation. Then, REMS-R lasted from ZT1-ZT9 and 20 min optogenetic stimulation was applied during NREM sleep, as described above (optogenetic protocols).

#### Gentle sleep deprivation (SD)

Animals were kept awake from ZT0-ZT4 while in their home cage and tethered to the recording apparatus. One person supervised two animals at the time and added nesting materials (pieces of Kleenex tissue) or gently interfered with the animals’ spontaneous behavior once it started adopting a sleeping posture^46^. For this, we used an ∼30 cm-long stick scotched to a ball of Kleenex tissue to gentle displace the litter around the nest or to wipe the wall of the cage. The animal was never touched throughout the entire procedure.

#### Stimulus-enriched sleep deprivation (SSD)

The animals were placed in a novel cage and subjected to several manipulations every 20-30 min to touching (1-2 min), gentle cage shaking (5 min) or auditory stimulation (knocking at the cage, 5 min) from ZT0-ZT4. Two animals were supervised per person. These manipulations were paired one each for touching and cage shaking and auditory stimulation and cage shaking, and once done the three together. At the end, animals were transferred back to their home cage and left undisturbed. In animals used for fiber photometric recording of LC fluorescence, one SSD was carried out. In animals in which LC was inhibited after SSD, two SSDs per animal were carried out (one for sham LED-off inhibition, one for light inhibition). Corticosterone measures were carried out at ZT4 for baseline (undisturbed) and SSD conditions per animal by drawing tail vein blood samples. Samples were kept on ice in heparin-coated tubes and centrifuged at 4° for 4 min 9,400xg. Plasma was extracted and corticosterone levels measured with the Enzo Life Sciences kit (Catalog No. ADI-901-097).

### HISTOLOGY

At the end of all the recording sessions the animal was injected i.p. with a lethal dose of pentobarbital. The position of each LFP electrode was subsequently marked using electro-coagulations (50 μA, 8 – 10 s). Afterwards, ∼40 mL paraformaldehyde (4%) were perfused intracardially at a rate of ∼3 mL min^-1^. The brains were then extracted and post-fixed for at least 24 h in 4% PFA at 4°C. Subsequently, the tissue was sliced in 50 μm-thick (for brainstem) or 100 μm-thick (for thalamus or cortex) sections using either a vibratome (Microtome Leica VT1000 S; speed: 0.25 – 0.5 mm s^-1^ and knife sectioning frequency: 65 Hz) or a freezing microtome (Microm). After cutting, the slices were either directly mounted (using Mowiol as a mounting medium) or stored in 0.1 M PB. Finally, we confirmed the anatomical location of the LFP and the viral-mediated expression of the fluorescent proteins with a Nikon SMZ25 Stereomicroscope equipped with a Nikon DS-Ri2 16 Mpx color camera. For a higher magnification image an Axiovision Imager Z1 (Zeiss) microscope was used (objectives used EC-Plan Neofluar 2.5x/0.075 ∞/0.17, 5x/0.16 ∞/0.17, 10x/0.3 ∞/-or 20x/0.5 ∞/0.17).

For assessing the colocalization of jGCaMP8s expression with tyrosine hydroxylase (TH) expression in the LC neurons, coronal brain sections 50 μm-thick (∼-5.3 mm from bregma), were first washed in 0.3% Triton in PBS 3 times, incubated in blocking solution of PBS 0.3 % and 2 % normal goat serum for 1 h and then overnight with the primary antibody (mouse anti-TH antibody, dilution 1:2000, Immunostar, 22941) at 4 °C while on a shaker. After a total incubation for at least 12 h, the slices were washed 3 times with PBS containing 0.3 % Triton. Afterwards, the secondary antibody was added (1:100 donkey anti-mouse antibody coupled to Alexa Fluor 594, Invitrogen, A21203, in PBS containing 0.3 % Triton) and the slices were incubated for 1-1.5 h at room temperature on a shaking platform. At the end of the procedure the slices were rinsed with PB 0.1 M and mounted (using again Mowiol as a mounting solution). The images (red and green emission channels) were acquired using a confocal microscope (Leica Stellaris 8) equipped with a 63x oil objective (HC PL APO 63x/1.40 oil CS2).

### ANALYSIS PROCEDURES

#### Sleep analysis

##### Sleep scoring

We scored sleep stages in 4-s epoch using the EEG/EMG signals according to standard procedures using a custom-made semiautomated Matlab routine available on our Github Repository (https://github.com/Romain2-5/IntanLuthiLab). For every epoch, EEG was inspected for its spectral composition and the EMG for its level of activity. Wakefulness was scored when high-frequency phasic EEG activity coincided with strong EMG activity; NREMS was scored when high-amplitude low-frequency components, such as slow oscillations (0.5 – 1.5 Hz) and δ activity (1.5 – 4 Hz), and recurrent sleep spindles (10 – 15 Hz) appeared in combination with low muscle tone; REMS was scored whenever a θ peak (6 – 10 Hz) appeared and muscle atonia developed. For determining the exact onset of REMS, a thresholding of θ/δ power was used for the CA1 LFP signal (see Spectral dynamics below). For epochs at transitions, the vigilance state that covered more than half of the epoch (> 2 s) was scored. Microarousals were scored when maximally 3 wake epochs (≤ 12 s) showing both EEG desynchronization (predominance of faster, low-amplitude, over slower high-amplitude activity) and EMG activity appeared sequentially and were both preceded and followed by NREMS. An epoch was also scored as MA when wake activity (in both EEG and EMG) lasted < 2 s. For analysis of intervals between successive REMS episodes during REMS-R, the motor vibration signal was used. The interval between two successive REMS episodes was quantified at the time spent in NREM sleep (in seconds) between two successive motor onsets (for this analysis microarousals were considered as part of NREM sleep).

##### Spectral dynamics

The dynamics of spectral bands were calculated from the S1 and CA1 LFPs using wavelet transforms with Gabor-Morlet Kernels with 4 cycles of standard deviation for the Gaussian envelope at 0.1 Hz resolution. For spectral analysis involving S1 LFP signals or EEG, frequency bands analyzed were δ (1.5 – 4 Hz), θ (6 – 10 Hz), σ (10 – 15 Hz), β (16 – 25 Hz), γ (60 – 80 Hz). The hippocampal LFP was used for calculation of the ratio of a θ (6 – 10 Hz) over δ (1 – 4 Hz) ratio (θ/δ) in the CA1 LFP electrode, in which the δ band was started at 1 Hz for convenience. For band-defined dynamic changes, the S1 or EEG signal were first down-sampled from 1 kHz to 200 Hz, the Wavelet transform calculated within the desired frequency bands and collapsing the frequency dimension. The derived signals were down-sampled at 10 Hz and low-pass-filtered at a cut-off frequency of 0.1 Hz. The detection of individual spindles was calculated as previously described^17,69^. For the CA1 signal, a similar procedure was applied to obtain the ratio for θ/δ and a threshold of 1.5 was chosen to define REMS onset around the previously defined manual transition. The dynamics of the EMG were calculated as absolute values from the 0.1 Hz-filtered trace.

##### Heart rate quantification

Heart rate was quantified as described before using the R-R peaks recorded by the EMG, using the square of its derivative and the Matlab FindPeaks function^17,21^.

#### Fiber photometric analysis

##### Extraction of ΔF/F_0_ signals and baseline correction

Using the modulated recorded photometry signal, we demodulated two separate channels. A functional signal (from the blue excitation LED), carrying the calcium-dependent activity, and an isosbestic, calcium-independent one (from the violet excitation LED). We utilized the workflow and the MATLAB codes available as open source^70^. Afterwards, the ΔF/F_0_ signal was computed using as F_0_ the fitted isosbestic signal, following the procedure explained in ref^71^.

For all analysis, fiber photometric signals were included in the analysis only if the dynamic range, calculated between the minimum and the maximum of the signal within the first 2 h of the recording, was > 15%. In the next paragraphs describing specific analyses, this extracted signal is referred to as the ‘LC signal’.

##### All-points histograms

We constructed all-points histograms from z-scored fiber photometric datasets obtained from all 10 animals for the first 2 h of the light phase. Overlap between NREMS and Wake period was calculated based on the number of data points in NREMS that are larger than the mean minus 2 standard deviations of the levels in wakefulness.

##### Analysis of LC peaks and LC activity transients

The definition of the LC peaks and activity transients are illustrated in **Extended Data Fig. 2** and are briefly summarized here. First, the LC signal was low-pass-filtered at 0.5 Hz (FIR filter of order 100) and rectified by the lower envelope (96-s sliding window) of the 0.1 Hz low-pass-filtered signal. Second, we detected the peaks in the rectified signal using the FindPeaks function of Matlab®. We discarded all peaks within the 60 percentile of prominence and used “halfheight” for the “WidthReference” parameter. From the rest of the peaks, we discarded the peaks under 25% of the maximum prominence and 20% of the signal amplitude. All MAs were included in these analyses.

Second, the LC activity transients in NREMS were defined as the fiber photometric signal events that show a rapid onset and that could contain multiple peaks. These were identified from the LC peaks within consolidated NREM sleep (>96 s of NREM sleep and after 1.5 times the time constant (Tau) of lower envelope decay after wake periods of >60 s). The maximum peak within a window of 20 s was defined as the transient peak. To define the start and the end of the transients we used the z-score of the 100 s window around each transient peak. The start of the transient was defined as the first peak within the derivate of the signals within the 10 s period before the transient peak. The end of the transient was the moment when the signal reached 15% of the maximal value of the 100 s window. The start and the onset of each activity transient were used to determine the duration and the area of the activity transients.

For the analysis of LC peaks after SD and SSD, mean peak frequency values were calculated for equal times spent in NREMS (12 bins for baseline conditions, 8 for the SD and SSD conditions).

##### Cross-correlations between LC activity, NA levels, σ power

Cross-correlations were calculated analogously to the previous one done for NA fluctuations and σ power^17^ using z-scored σ power and LC signals, or the NA and LC signals, at 0.1-s resolution, for all NREMS bouts > 96 s, including MAs, from ZT0-ZT12. A mean cross-correlation per animal was calculated and presented for the time interval between −200 s to 200 s. In the case of the cross-correlations between LC activity and σ power, the position of side peaks could be extracted in 6 out of 8 cases (based on the dynamic range of the LC signal, see **Extended Data Fig. 1e2**). For the cross-correlation between NA and LC signals (**Extended Data Fig. 1d**), the two animals with dually recorded fiber photometry were used. Cross-correlations were then averaged across animals. A positive time lag indicates that the LC signal increases preceded σ power decreases or NA increases. We refer to ‘LC activity transients’ as the high LC activity levels that anticorrelate with the σ power and that appear on infraslow time intervals (30 – 50 s).

##### Spectral analysis around LC activity peaks

Local maxima of the LC signals were defined using a 20-s sliding window on the LC ΔF/F_0_ signal, excluding the ones falling within the initial decay of the fiber photometric signal at transitions from Wake to NREMS. We then derived the z-scored spectral dynamics for different frequency bands using the wavelet approached previously described (See *Sleep Analysis - Spectral dynamics*) around these maxima from – 150 s to + 50 s. In a second step, these peaks were classified according to whether or not they were associated with a MA (defined by EEG/EMG activity as described in *Sleep Analysis – Sleep Scoring*) within 5 s around the peak. Additionally, we calculated the absolute EMG activity and the z-scored heart rate within these windows and repeated this analysis for both for the S1 LFP and the EEG during NREMS bouts for all times. Mean amplitudes for the LC signal and for heart rate were calculated as the difference from the peak time to the minimum value before the peak (−10 s to 0 s). Mean amplitudes for spectral power bands were calculated from times 0 s to 1 s.

##### Spectral analysis at REMS onset

The exact timing of REMS onset was defined using a thresholding for θ/δ at 1.5 using the CA1 LFP electrode. Using the parietal EEG electrode provided similar time points. The dynamics for preceding NREMS were then obtained for the S1 LFP σ band and for the low-pass filtered absolute EMG signal. For LC signals with high dynamic range, an inflection point could be observed prior to REMS onset after which there was a further decrease to a new minimum (see e.g. Fig. 2c). This inflection point was associated with a further increase in σ power and the onset of EMG decline, consistent with markers for ‘Intermediate sleep’.

##### Sleep architecture analysis during LC optogenetic stimulation and inhibition

For LC stimulation or inhibition according to the phase of the σ power fluctuation, times spent in Wake, NREMS and REMS were calculated for the test or sham stimulation 20-min sessions and expressed in % of total time. Transitions were calculated as the number of REMS-epochs followed by NREMS. For LC inhibition at different moments in the NREMS-REMS cycle, a similar procedure was applied.

##### Mean LC activity in NREMS bouts

The LC fluorescent signal was extracted between REMS bouts lasting ≥ 12 s, z-scored for all NREMS including MAs, and plotted over normalized NREM time between the consecutive REM bouts. The derived signals were averaged based on the length of the preceding REM bout as previously done for similar purposes^44^.

##### Quantification of rebound REMS after REMS-R

To demonstrate the efficiency of the REMS-R, we also quantified rebound REMS by calculating cumulative times spent in REMS during REMS-R and the 6 h – post REMS-R period.

##### LC signal analysis during REMS-R

To determine the relative level of LC activity prior to REMS entry during the REMS-R, we extracted bouts where a transition to REMS was present and preceded by at least 12 s of NREM sleep. The ΔF/F_0_ LC activity was then z-scored for each bout and we reported the mean ΔF/F_0_ value during the last 5 s before the transition to REMS.

##### LC stimulation during REMS-R

For each 20 min stimulation block (first 20 min of every h from ZT1– ZT9) during the REMS-R, we normalized the time in equal amounts of NREMS for the periods before, during and after stimulation (10 bins per block) and calculated the number of REM entries per min of NREMS. A similar approach was done to the analogous periods during sham stimulation, *ceteris paribus*, with the LEDs turned off.

### STATISTICS

The statistics were done using Python, Matlab R2021a or R statistical language version 4.0.1. The normality of data distributions was assessed using the Shapiro–Wilk and Levene tests. In cases of normality violations, non-parametric Kruskal-Wallis and Wilcoxon tests were used. Datasets were collected in experimental designs allowing preferentially repeated measures and/or paired statistical analysis. Post-hoc analyses were done only when the one-way factor or the interaction between factors was significant (p<0.05). Bonferroni’s correction for multiple comparisons was applied routinely, and the corrected α thresholds are given in the figure legends. Differences in the dynamics of power bands and heart rate were calculated using t-tests and correcting for multiple comparisons using False Discovery Rates (FDR) where the significant values satisfy P(*k)< 0.05* k/2000,* where P is the vector of the ranked p values of the point-to-poitn t-test within the window (−150 s to 50 s at 10 Hz). In all figure legends, the statistical tests used are mentioned, while sample numbers and p values are indicated directly in the figure panels. Details of all statistical tests and effect sizes are summarized in **Supplementary Table 1**.

