## Supplementary material for "Noradrenergic *locus coeruleus* activity functionally partitions NREM sleep to gatekeep the NREM-REM sleep cycle": Suppl Figures 1-6 and Suppl Table 1

### functionally partitions NREM sleep to

#### Extended Data Figures and Supplementary Table.

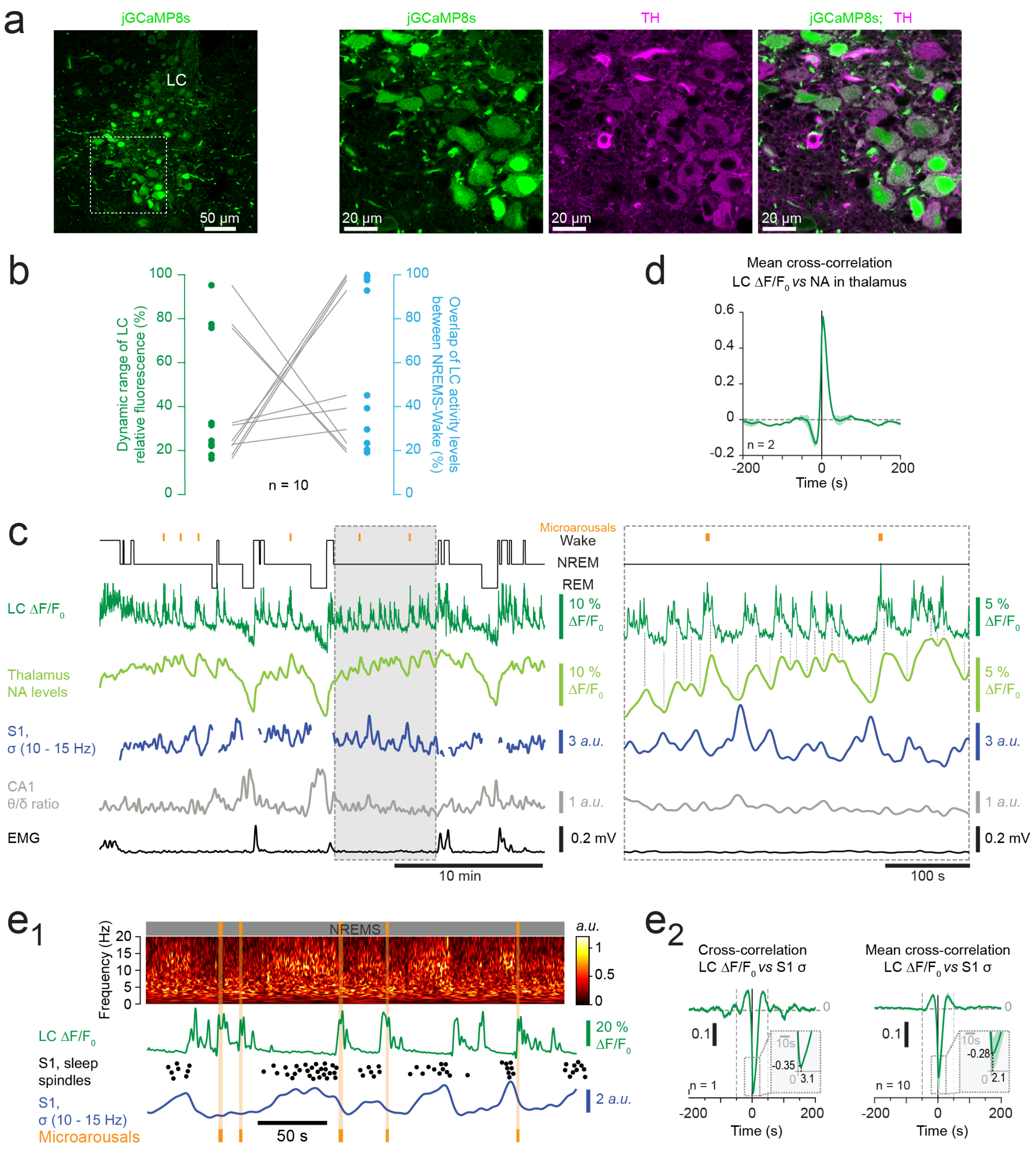


**Extended Data Figure 1. Histological verification of viral expression specificity and additional analyses.**

a) Confocal micrographs taken from a representative DBH-Cre mouse expressing jGCaMP8s (green, same animal as **Fig. 1c**) in the LC. Immunostaining for tyrosine hydroxylase (TH, violet) shows the specificity of viral targeting, evident by overlap of green and violet fluorescence within LC neuronal somata. Dotted square in the left image indicates the area expanded on the right.

b) Scatter plot highlighting the relationship between the dynamic range of the LC fluorescence signal (green dots) and the overlap of LC activity levels between Wake and NREMS (blue dots), as calculated up to a a threshold of mean - 2 standard deviations of wake levels for n = 10 animals included in the analysis of Figure 1. High dynamic ranges show the lowest overlap, whereas low dynamic ranges provide more variable values of overlap. The mean value of overlap indicated in the main text was calculated by taking the average across the blue datapoints.

c) Example dual fiber photometric recording from an animal that expresses jGCaMP8s in LC and $GRAB_{NE1h}$ in somatosensory thalamus. From the top, Hypnogram (black), LC-jGCaMP8s and free NA-$GRAB_{NE1h}$ fluorescence, S1 LFP sigma (σ, 10 – 15 Hz) power, CA1 local field potential (LFP) theta/delta (θ/δ) ratio and absolute EMG levels. Portion of trace underlain in grey expanded on the right. Vertical lines highlight the coordination between LC and NA fluorescence signals.

d) Mean cross-correlation of LC activity with free NA levels for 2 dual fiber photometric recordings in two DBH-Cre animals expressing GCaMP8s in LC and the NA sensor $GRAB_{NE1h}$ in primary somatosensory thalamus. Data from one such recording are shown in panel c. Cross-correlation coefficients were 0.56 and 0.59 at a positive lag of 2.9 and 3.1 s for the two recordings.

e) Anticorrelation of LC activity and σ power. e1) Time-frequency plot of S1 LFP in NREMS, aligned with LC fluorescent signal. Data from the recording shown in Fig. 1c. S1 LFP σ (10 – 15 Hz) power and individual sleep spindles (black dots) shown below. Vertical orange lines: MAs; e2) Cross-correlations for LC fluorescence and S1 σ power from ZT0-ZT12 (same animals as e1) and mean cross-correlation across n = 10 animals. Vertical dotted lines: side peaks. Insets; zoom-ins with crosscorrelation magnitude and lag values.

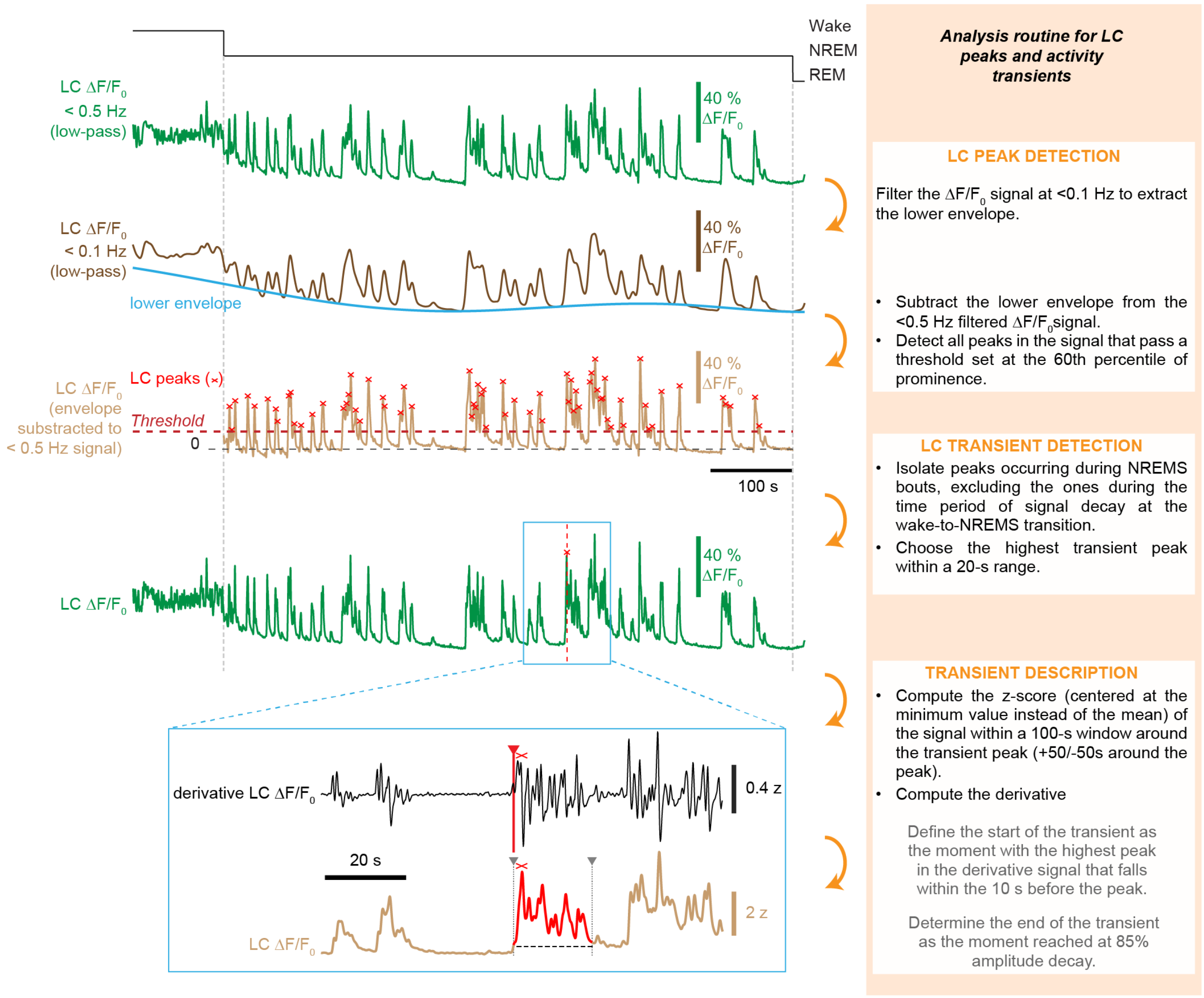


**Extended Data Figure 2. Illustration of analysis routine to identify LC activity peaks and transients.**

Step-by-step illustration of the algorithm used, starting from the original LC fluorescence signals. From top to bottom, traces shown are: Hypnogram (black), corresponding LC fluorescent signal (green), low-pass-(0.1 Hz) filtered signal (dark brown) with lower envelope (blue), subtracted trace (light brown). On this last signal, peak analysis was carried out using a threshold and peak prominence >40% (for more details, see Methods). Detected peaks are indicated by red crosses. From these individual peaks, activity transients are identified, of which an example is given in the expanded trace portion surrounded by a blue rectangle.

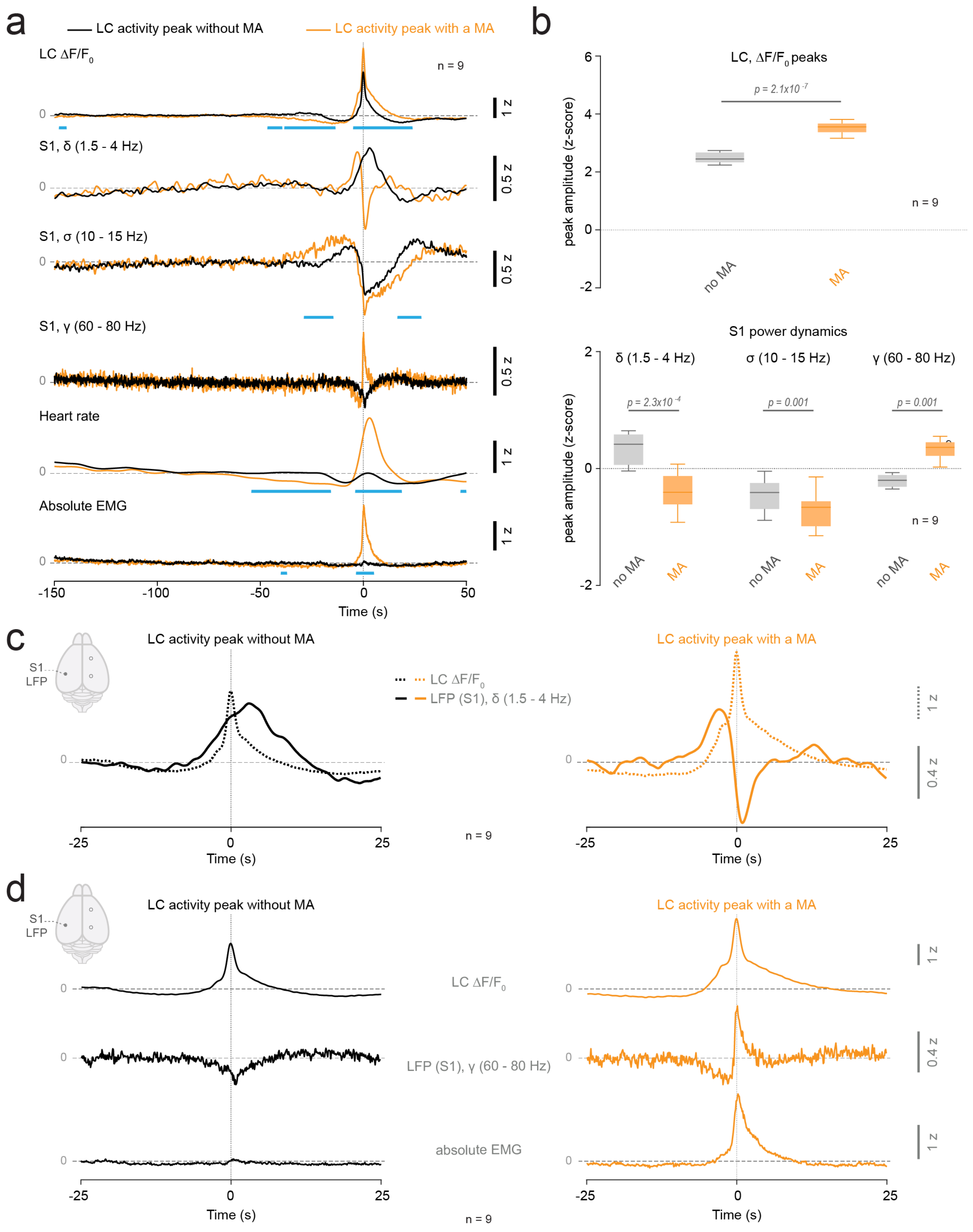


**Extended Data Figure 3. Extended spectral analysis of LC activity transients**

a) Spectral analysis of S1 LFP signals corresponding to LC fluorescence transients that are classified based on whether they did not (black traces) or did co-occur with a MA (orange traces). Corresponding power dynamics for the delta (δ), sigma (σ) and gamma (γ) frequency bands are aligned vertically, together with heart rate and absolute EMG. Traces are means across n = 10 animals. Blue bars denote significance as calculated by false discovery rates.

b) Quantification of mean values between 0 – 5 s. Statistical analysis through paired t-tests, with Bonferroni corrected p for power bands = 0.017.

c) Overlay of LC fluorescence (dotted line) and $\delta$ power (continuous line) on an expanded time scale, with traces taken from panel a, with the same color code. Note the increase in $\delta$ power accompanying the onset of the LC activity peak that becomes inverted to reach negative values in the case of a MA-associated LC activity peak (orange trace).

d) As c, for LC fluorescence and $\gamma$ power.

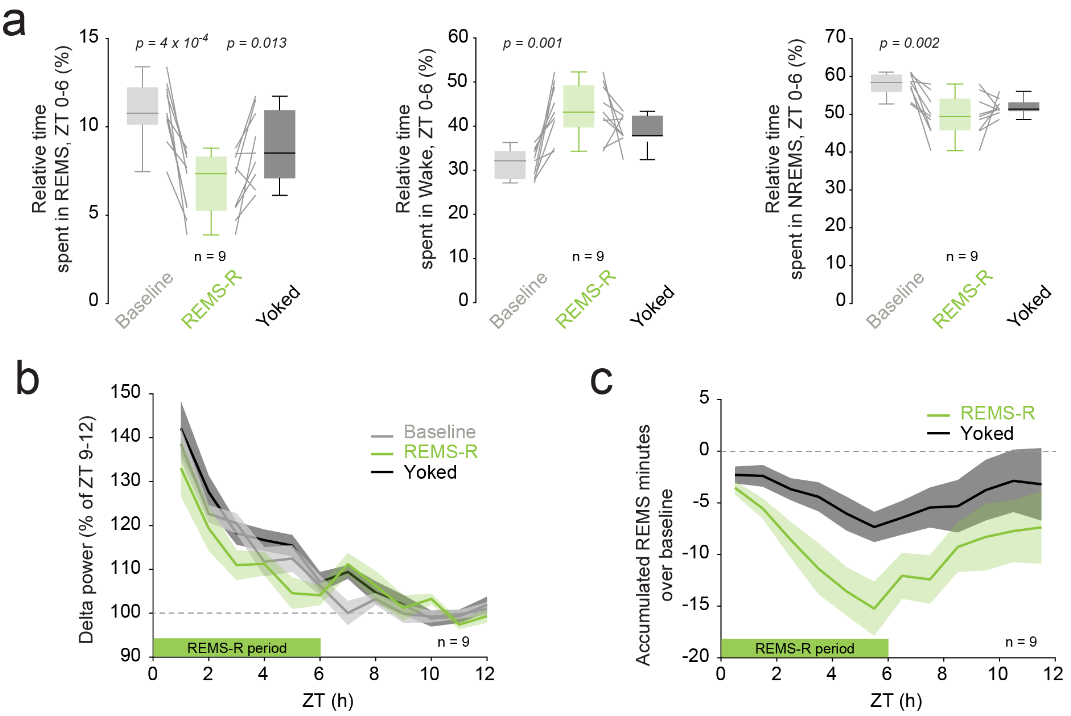


**Extended Data Figure 4. Validation of the efficiency and specificity of the REMS-R technique.**

a) Box-and-whisker plots of times spent in REMS, wake and NREMS for baseline, REMS-R and yoked conditions. Experiment was done in a paired design, with each animal once used for REMS-R and as yoked control. Paired data are connected with grey lines. Paired t-tests with Bonferroni correction.

b) The decline in low-frequency $\delta$ power (1.5 – 4 Hz) power in the 12-h light phase during which REMS-R was carried out from ZT0-ZT6.

c) Cumulated time spent in REMS during the REMS-R and the subsequent recovery period in undisturbed conditions. Note the greater loss of REMS time during REMS-R compared to yoked animals, but recovered afterwards.

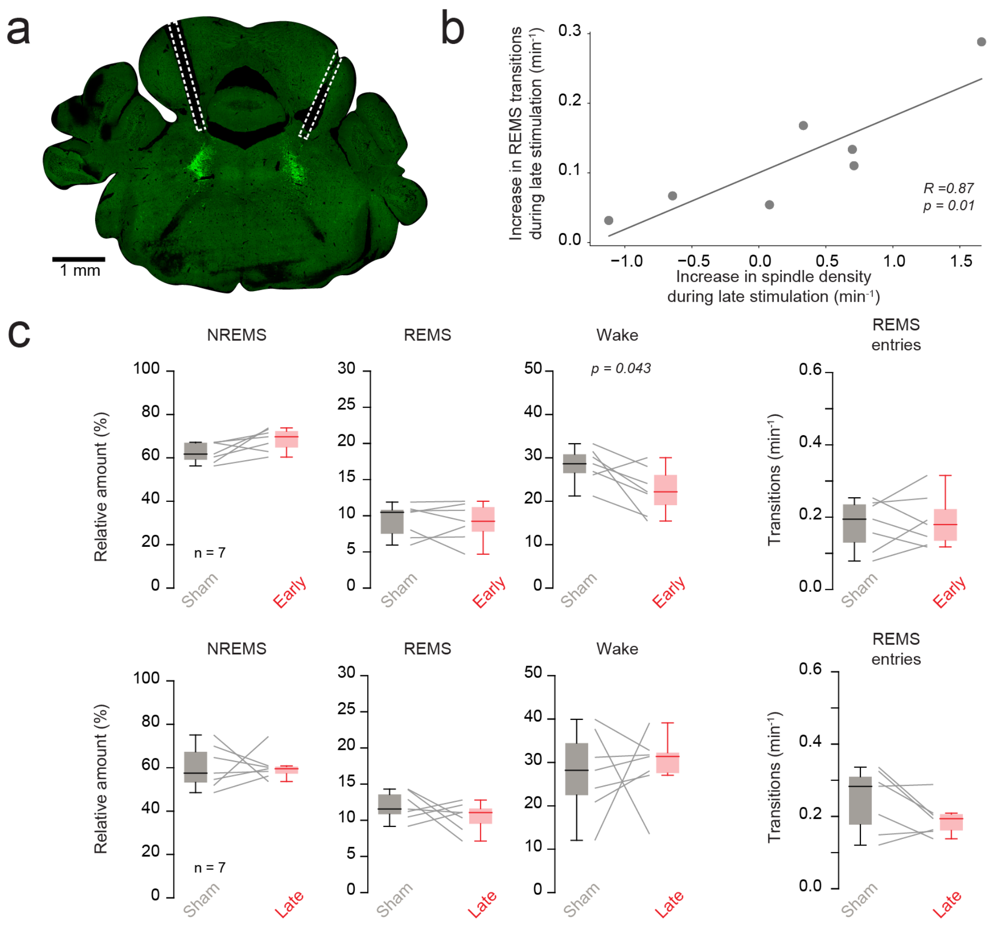


**Extended Data Figure 5. Validating the efficiency of Jaws-mediated inhibition of LC using histological and functional methods.**

a) Example fluorescent micrograph of a mouse illustrating EGFP fluorescence of Jaws-expressing LC neurons and position of optic fibers on top of LC. Wilcoxon signed-rank test.

b) Linear correlation between light-induced sleep spindle density changes and REMS transitions for the experiment involving LC inhibition late in the undisturbed NREMS-REMS cycle (Fig. 4d).

c) Summary data of control experiments for the experiments described in Figure 4 using animals expressing non-light-sensitive (mCherry-expressing) viral constructs. Sham corresponds to LED-off conditions. Paired t-tests or Wilcoxon signed-rank tests.

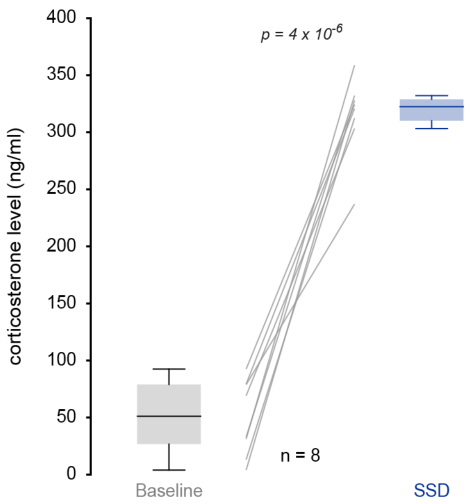


**Extended Data Figure 6. Plasma corticosterone levels after SSD in comparison to undisturbed animals at the same time of day.**

Box-and-whisker plots, with grey lines connecting paired datasets taken at ZT4, once in baseline undisturbed sleep, once after a 4-h SSD. Paired t-test.

**Supplementary Table S1. Statistical table for Main and Extended Data Figures.**

This table describes statistical tests used for each figure/panel in this paper. Per figure panel, the number of animals, the test statistics (F/t values) and the degrees of freedom (Df/df) are indicated. P values are given in a separate column. Bonferroni-corrected p values are given in the legends when appropriate. Effect size is given whenever significance was reached.

| Figs and panels | Test used | n numbers (animals unless otherwise indicated) | Test statistics (F, t) & degrees of freedom (Df, df) | P Value | *Post hoc* tests with multiple comparisons, Test statistics & df | *Post-hoc tests*, p values) | Effect size (Cohen's D) |
|---|---|---|---|---|---|---|---|
| **1d** | One-way RM ANOVA with factor 'vigilance state) | 10 | F= 472.3, Df = 9 | 4.36 * $10^{-9}$ | **For Wake vs NREMS**: t = 6.192, df = 9<br><br>**For NREMS vs REMS**: t = 4.641, df = 9 | **For Wake vs NREMS**: p=0.00016<br><br>**For NREMS vs REMS**: p = 0.0012 | **For Wake vs NREMS**: D=3.24<br><br>**For NREMS vs REMS**: D=2.08 |
| **1g** | Student's paired t-test | 10 | **For duration of activity transients**: t = -2.679; Df = 9<br><br>**For area of activity transient**: t = -6.959; Df = 9 | **For duration of activity transients**: p = 0.025<br><br>**For area of activity transients**: p = 6.6 * $10^{-5}$ | | | **For duration of activity transient:** D = -0.48<br><br>**For area of activity transient**: D = -1.11 |
| **1i** | Student's paired t-test for measures between non-MA-associated and | 10 | **For $\Delta F/F_0$ signal**: t = -15.04, Df = 9<br><br>**For delta power**: | **For $\Delta F/F_0$ signal**: p = 1.09*$10^{-7}$<br><br>**For delta power**: | | | **For $\Delta F/F_0$ signal**: D = -6.77<br><br>**For delta power:** D = 3.95 |

| | | | | | | | |
|---|---|---|---|---|---|---|---|
| | MA-conditions | | t = 10.21, Df = 9<br>**For sigma power**:<br>t = 9.74, Df=9<br><br>**For gamma power**:<br>t=-13.09, Df=9<br><br>**For heart rate**:<br>t = -9.67, Df=9 | p = $3.02*10^{-6}$<br>**For sigma power**:<br>p = $4.4*10^{-6}$<br><br>**For gamma power**:<br>p = $3.66*10^{-7}$<br><br>**For heart rate**:<br>p = $4.73*10^{-6}$ | | | **For sigma power:**<br>D = 2.54<br><br>**For gamma power**:<br>D = -4.71<br><br>**For heart rate**:<br>D = -3.87 |
| **2b, left** | Wilcoxon signed-rank for effects of LC stim compared to Sham Stim on % Time spent in Wake, NREMS and REMS and on REMS entries | 9 | **For Wake**: V = 28<br><br>**For NREMS**: V = 4<br><br>**For REMS**: V=44<br><br>**For REMS entries**: V=44 | **For Wake**: p = 0.6<br><br>**For NREMS**: p = 0.03<br><br>**For REMS**: p=0.008<br><br>**For REMS entries**: p=0.008 | | | **For NREMS**: D = -1.28<br><br>**For REMS**: D = 1.97<br><br>**For REMS entries**: D = 2.28 |
| **2b, right** | Wilcoxon signed-rank for effects of LC inhibition compared to Sham Stimulation on % Time spent in Wake, NREMS and REMS | 9 | **For Wake**: V = 43<br><br>**For NREMS**: V = 22<br><br>**For REMS**: V=1<br><br>**For REMS entries**: V=0 | **For Wake**: p = 0.13<br><br>**For NREMS**: p 0.63<br><br>**For REMS**: p=0.004<br><br>**For REMS entries**: p=0.002 | | | **For REMS**: D = -1.35<br><br>**For REMS entries**: D = -1.08 |

| | | | | | | | |
|---|---|---|---|---|---|---|---|
| **2c** | Wilcoxon signed-rank tests | N = 45 events for n = 2 animals | Test statistics for Wilcoxon test: V=269.0 | p = 0.0043 | | | |
| **2e** | One-way RM ANOVA with factor 'moment in NREMS bout' | 10 | F = 8.54, Df = 1 | p = 0.017 | For time points 0.3-0.4 vs 0.8-0.9 of NREMS bout: t = -3.0216, df = 9 | p = 0.4936 | |
| **3b** | **Transitions to REMS:** Wilcoxon signed-rank | 9 | **Baseline vs REMSD:** V =0<br><br>**REMSD vs Yoked:** V = 45 | **Baseline vs REMSD:** p = 0.0039<br><br>**REMSD vs Yoked:** p = 0.0039 | | | **Baseline vs REMSD:** D = -2.04<br><br>**REMSD vs Yoked:** D = 1.91 |
| **3e1,e2** | **Inter REMS interval:** Paired t-test<br><br>**LC activity before REMS entry:** Paired t-test | 8 | **Inter REMS interval:** t = 5.86; df = 7<br><br>**LC activity before REMS entry:** t =-0.698; df = 7 | **Inter REMS interval:** p = 0.0006<br><br>**LC activity before REMS entry:** p = 0.514 | | | **Inter REMS interval:** D = 1.93 |
| **3f** | Wilcoxon signed-rank | 6 | V=0 | p = 0.031 | | | D = 3.16 |
| **4c** | **Transitions to REMS:** Wilcoxon signed-rank<br><br>**Time spent in** | 7 | **Transitions to REMS:** V = 10.0<br><br>**Time spent in** | **Transitions to REMS:** p = 0.58<br><br>**Time spent in REMS:** | | | |

| | | | | | | | |
|---|---|---|---|---|---|---|---|
| | **REMS**: Paired t-test<br><br>**Time spent in NREMS**: Paired t-test<br><br>**Time spent in Wake**: Paired t-test | | **REMS:** t = -0.7263; df = 6<br><br>**Time spent in NREMS:** t = 7.3184; df = 6<br><br>**Time spent in Wake**: t = -5.1531; df = 6 | p = 0.4950<br><br>**Time spent in NREMS:** p = 0.0003<br><br>**Time spent in Wake:** p = 0.0021 | | | <br><br>**For time spent in NREMS**: D = 2.28<br><br>**For time spent in wake**: D = -2.59 |
| **4d** | **Transitions to REMS:** Wilcoxon signed-rank<br><br>**Time spent in REMS**: Paired t-test<br><br>**Time spent in NREMS**: Paired t-test<br><br>**Time spent in Wake**: Paired t-test | 7 | **Transitions to REMS**: V = 0.0<br><br>**Time spent in REMS:** t = -3.6183; df = 6<br><br>**Time spent in NREMS:** t = 2.8396; df = 6<br><br>**Time spent in Wake**: t = -1.6369; df = 6 | **Transitions to REMS**: p = 0.0156<br><br>**Time spent in REMS**: p = 0.0111<br><br>**Time spent in NREMS:** p = 0.0295<br><br>**Time spent in Wake:** p = 0.1527 | | | **Transitions to REMS**: D = -1.76<br><br>**Time spent in REMS**: D = -0.80<br><br>**Time spent in NREMS:** D = 1.38 |
| **5c** | Paired t-test | 7 | t = -0.145753; df = 8 | p = 0.8877 | | | |

| | | | | | | | |
|---|---|---|---|---|---|---|---|
| **5d** | Paired t-test | 7 | t = 6.9188; df = 8 | p = 0.000122 | | | D = -2.86 |
| **5e** | Paired t-test | 7 | t = 3.84; df = 6 | p = 0.008 | | | D = -184 |
| **5f** | **REMS Latency:** Wilcoxon signed-rank<br><br>**Transitions to REMS:** Paired t-test | 9 | **REMS Latency:** V = 1.0<br><br>**Transitions to REMS:** t = 3.2497; df = 8 | **REMS Latency:** p = 0.007<br><br>**Transitions to REMS:** p = 0.011 | | | D = -1.59<br><br>D = 1.5833 |
| **6a** | Student's paired t-test for measures between non-MA-associated and MA-conditions | 7 | **$\Delta F/F_0$ signal**: t = -8.2717, df = 6<br><br>**Delta power**: t = 7.1402, df = 6<br><br>**Sigma power**: t = 4.9354, df = 6<br><br>**Gamma power**: t = -8.2717, df = 6 | **$\Delta F/F_0$ signal**: p = 0.00017<br><br>**Delta power**: p = 0.00038<br><br>**Sigma power**: p = 0.0026<br><br>**Gamma power**: p = 0.00044 | | | **For $\Delta F/F_0$ signal**: D = -3.48<br><br>**For Delta power**: D = 3.31<br><br>**For Sigma power**: D = 2.12<br><br>**For Gamma power**: D = -4.49 |
| **6c** | **MA density:** Paired t-test<br><br>**REM latency:** | 9 | **MA density:** t = 2.677; df = 8<br><br>**REM latency:** | **MA density:** p = 0.0280<br><br>**REM latency:** p = 0.0149 | | | **For MA density:** D = 0.85<br><br>**For REM latency:** |

| | | | | | | | |
|---|---|---|---|---|---|---|---|
| | Paired t-test | | t = 3.086<br>df = 8 | | | | D = 1.36 |
| **Extended Data Fig 3b** | Student's paired t-test for measures between non-MA-associated and MA-conditions | 9 | **$\Delta F/F_0$ signal**: t = -16.251, df = 8<br><br>**Delta power**: t = 6.3263, df = 8<br><br>**Sigma power**: t = 4.972, df = 8<br><br>**Gamma power**: t = -4.8073, df = 8 | **$\Delta F/F_0$ signal**: p = $2.07*10^{-7}$<br><br>**Delta power**: p = 0.00023<br><br>**Sigma power**: p = 0.0011<br><br>**Gamma power**: p = 0.0013 | | | **For $\Delta F/F_0$ signal**: D = -5.17<br><br>**For Delta power**: D = 2.48<br><br>**For Sigma power**: D = 0.88<br><br>**For Gamma power**: D = -2.25 |
| **Extended Data Fig 4a** | **Time spent in REMS:** Paired t-test<br><br>**Time spent in Wake:** Paired t-test | n = 9 | **Time spent in REMS:** For Baseline vs REMS-R: t = 5.822; df = 8<br>For REMS-R vs yoked: t = -3.171; df = 8<br><br>**Time spent in Wake:** For Baseline vs REMS-R: t = -4.933; df = 8<br><br>For REMS-R | **Time spent in REMS:** For Baseline vs REMS-R: p = 0.0004<br>For REMS-R vs yoked: p = 0.0132<br><br>**Time spent in Wake:** For Baseline vs REMS-R: p = 0.0011<br><br>For REMS-R vs yoked: p = 0.1459 | | | **Time spent in REMS:** D = 2.34<br>D = -1.11<br><br>**Time spent in Wake:** D = -2.41 |

| | | | | | | | |
|---|---|---|---|---|---|---|---|
| | **Time spent in NREMS:** Paired t-test | | vs yoked: t = 1.611; df = 8<br><br>**Time spent in NREMS:** For Baseline vs REMS-R: t = 4.346; df = 8<br><br>For REMS-R vs yoked: t = -1.024; df = 8 | **Time spent in NREMS:** For Baseline vs REMS-R: p = 0.0025<br><br>For REMS-R vs yoked: p = 0.3357 | | | **Time spent in NREMS:** D = 1.78 |
| **Extended Data Fig 5c** | <u>**Early**</u><br><br>**Transitions to REMS:** Paired t-test<br><br>**Time spent in REMS**: Paired t-test<br><br>**Time spent in NREMS**: Paired t-test<br><br>**Time spent in Wake**: Paired t-test | n = 7 | <u>**Early**</u><br><br>**Transitions to REMS:** t = -0.4087; df = 6<br><br>**Time spent in REMS**: t = 0.1955; df = 6<br><br>**Time spent in NREMS**: t = -2.3997; df = 6<br><br>**Time spent in Wake**: t = 2.5490; df = 6 | <u>**Early**</u><br><br>**Transitions to REMS**: p = 0.6968<br><br>**Time spent in REMS**: p = 0.8514<br><br>**Time spent in NREMS:** p = 0.053<br><br>**Time spent in Wake:** p = 0.0435 | | | |

| | | | | | | | |
|---|---|---|---|---|---|---|---|
| | **Late:**<br><br>**Transitions to REMS:** Paired t-test<br><br>**Time spent in REMS**: Paired t-test<br><br>**Time spent in NREMS**: Wilcoxon signed-rank<br><br>**Time spent in Wake**: Paired t-test | | **Late**<br><br>**Transitions to REMS**: t = 1.9133; df = 6<br><br>**Time spent in REMS:** t = 1.1423; df = 6<br><br>**Time spent in NREMS:** V = 13; df = 6<br><br>**Time spent in Wake**: t = -0.2941; df = 6 | **Late**<br><br>**Transitions to REMS**: p = 0.1042<br><br>**Time spent in REMS**: p = 0.2968<br><br>**Time spent in NREMS:** p = 0.9375<br><br>**Time spent in Wake:** p = 0.8226 | | | |
| **Extended Data Fig. 6** | t-test | 8 | t = -12.8; df = 7 | p = $4*10^{-6}$ | | | D = -7.65 |
